# Chromatin accessibility differences between alpha, beta, and delta cells identifies common and cell type-specific enhancers

**DOI:** 10.1101/2021.12.06.471006

**Authors:** Alex M. Mawla, Talitha van der Meulen, Mark O. Huising

**Affiliations:** Department of Neurobiology, Physiology & Behavior, College of Biological Sciences, University of California, Davis, CA 95616, USA; Department of Physiology and Membrane Biology, School of Medicine, University of California, Davis, CA 95616, USA

**Author notes:** Corresponding author: Dr. Mark O. Huising, Ph.D., Professor, University of California, Davis, Department of Neurobiology, Physiology & Behavior, College of Biological Sciences, Department of Physiology and Membrane Biology, School of Medicine, One Shields Avenue, Davis, CA, 95616.

## Abstract

High throughput sequencing has enabled the interrogation of the transcriptomic landscape of glucagon-secreting alpha cells, insulin-secreting beta cells, and somatostatin-secreting delta cells. These approaches have furthered our understanding of expression patterns that define healthy or diseased islet cell types and helped explicate some of the intricacies between major islet cell crosstalk and glucose regulation. All three endocrine cell types derive from a common pancreatic progenitor, yet alpha and beta cells have partially opposing functions, and delta cells modulate and control insulin and glucagon release. While gene signatures that define and maintain cellular identity have been widely explored, the underlying epigenetic components are incompletely characterized and understood. Chromatin accessibility and remodeling is a dynamic attribute that plays a critical role to determine and maintain cellular identity. Here, we compare and contrast the chromatin landscape between mouse alpha, beta, and delta cells using ATAC-Seq to evaluate the significant differences in chromatin accessibility. The similarities and differences in chromatin accessibility between these related islet endocrine cells help define their fate in support of their distinct functional roles. We identify patterns that suggest that both alpha and delta cells are poised, but repressed, from becoming beta-like. We also identify patterns in differentially enriched chromatin that have transcription factor motifs preferentially associated with different regions of the genome. Finally, we identify and visualize both novel and previously discovered common endocrine- and cell specific- enhancer regions across differentially enriched chromatin.

## Introduction

The evaluation of the transcriptional landscape through high-throughput bulk sequencing (bulkSeq) in both mouse and human of major islet cell types has granted a deeper understanding of cellular identity and intercellular crosstalk within the pancreas. This has enabled the detection of distinct gene pattern signatures between major islet cell types in mouse and human [1–6]. However, gene expression represents the final outcome of a complex layer of genetic and epigenetic factors that determine islet cell fate [7–9] and identity [10, 11]. Previous studies have explored pancreatic islet cellular identity by evaluating epigenomic features such as methylation [12–14], histone modifications [15–18], and enhancer regulatory regions [19–24]. While each of these factors contributes to defining and maintaining cell fate and identity, connecting chromatin accessibility differences to epigenetic factors promises to provide further insight into outstanding questions within the field.

Chromatin remodeling is a central epigenetic regulator that can be surveyed in order to better understand cell states [20, 25–28]. The accessibility of chromatin via changes between euchromatin and heterochromatin, and nucleosome occupancy, plays a significant role in cell lineage, and in tissue- and cell-specific gene expression [11, 25, 29]. Epigenetic stability is required for the maintenance of islet cell identity, while changes in chromatin accessibility are associated with perturbations in gene expression due to disease [7, 22, 30]. Chromatin accessibility in tandem with other epigenetic factors at promoter-proximal regions [29, 31] of a gene allows for direct activation or repression of transcription. In contrast, open chromatin at exonic [32], intronic [33], or distal-intergenic regions [34] can be accessed by regulatory factors that act as nearby or distal enhancers that govern lineage branching and stable cell fate.

Assay for transposase-accessible chromatin using sequencing (ATAC-Seq) allows for the unbiased, modification-independent evaluation of chromatin accessibility within cell types and can be run with relatively small sample input [30, 35]. Previous studies have explored chromatin accessibility in healthy [5, 11, 36, 37] and T2D [22, 23, 38, 39] islets as well as pancreatic progenitors [9] using bulkSeq through human antibody panels alongside FACS-purification or through single-cell sequencing (scATACSeq) [40–42]. However, none of these studies have explored pancreatic islet cell chromatin accessibility from mouse FACS-purified alpha, beta, and delta cells. Therefore, to better understand endocrine islet cell identity between mouse alpha, beta, and delta cells, we compared chromatin accessibility and transcriptome data for FACS-purified mouse alpha, beta, and delta cells sorted from pancreatic islets from triple transgenic reporter strains - mIns1-H2b-mCherry beta cells crossed to mice with alpha or delta cells marked by YFP in a Cre-dependent fashion - that we generated for this purpose [1, 6]. This approach allowed for the direct comparison between ATAC-Seq and RNA-Seq datasets from alpha, beta, and delta cells from these lines.

We integrated our ATAC-Seq data with high-quality transcription factor and histone binding data from other mouse pancreatic islet studies to evaluate how transcriptional activators and repressors may collectively regulate differential gene expression at promoter-proximal regions. To support the visualization and integration of our ATAC-Seq chromatin data and previously published transcriptome of the FACS-purified alpha, beta, and delta cells alongside select epigenomic datasets from histone marker and transcription factor Chromatin Immuno Precipitation (ChIP) data, we developed an R package, *epiRomics* [See: https://github.com/Huising-Lab/epiRomics]. This package is a novel, publicly available resource that is described in detail elsewhere [43]. *epiRomics* allows for the visualization of integrated epigenomic data and visualizes putative enhancer regions without the requirement for extensive bio-informatics experience, with the intent of enabling more of our colleagues to tease apart key regions that may drive cell fate switching and maintenance between the major islet endocrine cell types. Through this approach we identified putative enhancer regions at distal-intergenic regions common to all cell types as well as regions selectively accessible only in a single islet cell type and confirmed previously identified mouse pancreatic islet enhancers.

## Methods

### Islet isolation and FACS sorting

mIns1-H2b-mCherry [1] x Rosa-LSL-YFP crossed to either Sst-Cre [44] or Gcg-Cre [45] triple transgenic mice were pooled by sex, each sample yielding a median of 20,000 cells, with islet isolation and FACS-sorting as previously described (Supplemental Fig. 1) [1, 46–48].

### Assay for transposase-accessible chromatin using sequencing

Single-end 50 bp reads were generated after library size selection yielded an average of 450 bp fragments and sequenced as previously described using NexteraDNA library protocol [30].

### Alignment and differential peak calling

Reads from each replicate (Supplemental Table 1) were evaluated for quality control and trimmed using FastQC and Trimmomatic, respectively [49–51]. A modified index of Gencode GRCm38.p4 (mm10) was built to exclude mitochondrial DNA prior to aligning reads with Bowtie 2 [52, 53]. Post-alignment, duplicates were marked using Picard Tools, blacklist regions were removed, and BAM files were converted into tagAlign format for downstream use. Peak calling and bigwig generation was done using MACS2 [54]. Differential expression testing was performed using DiffBind’s edgeR method [55, 56].

### Quality control and validation

Quality control metrics were evaluated within raw reads as well as peak calls and compared against ENCODE standards for fraction of reads in peaks (FRiP), leading to the removal of one beta replicate with a FRiP score far below 0.3 (Supplemental Table 1; Fig. 1A-C) [57, 58].

**Figure 1.**
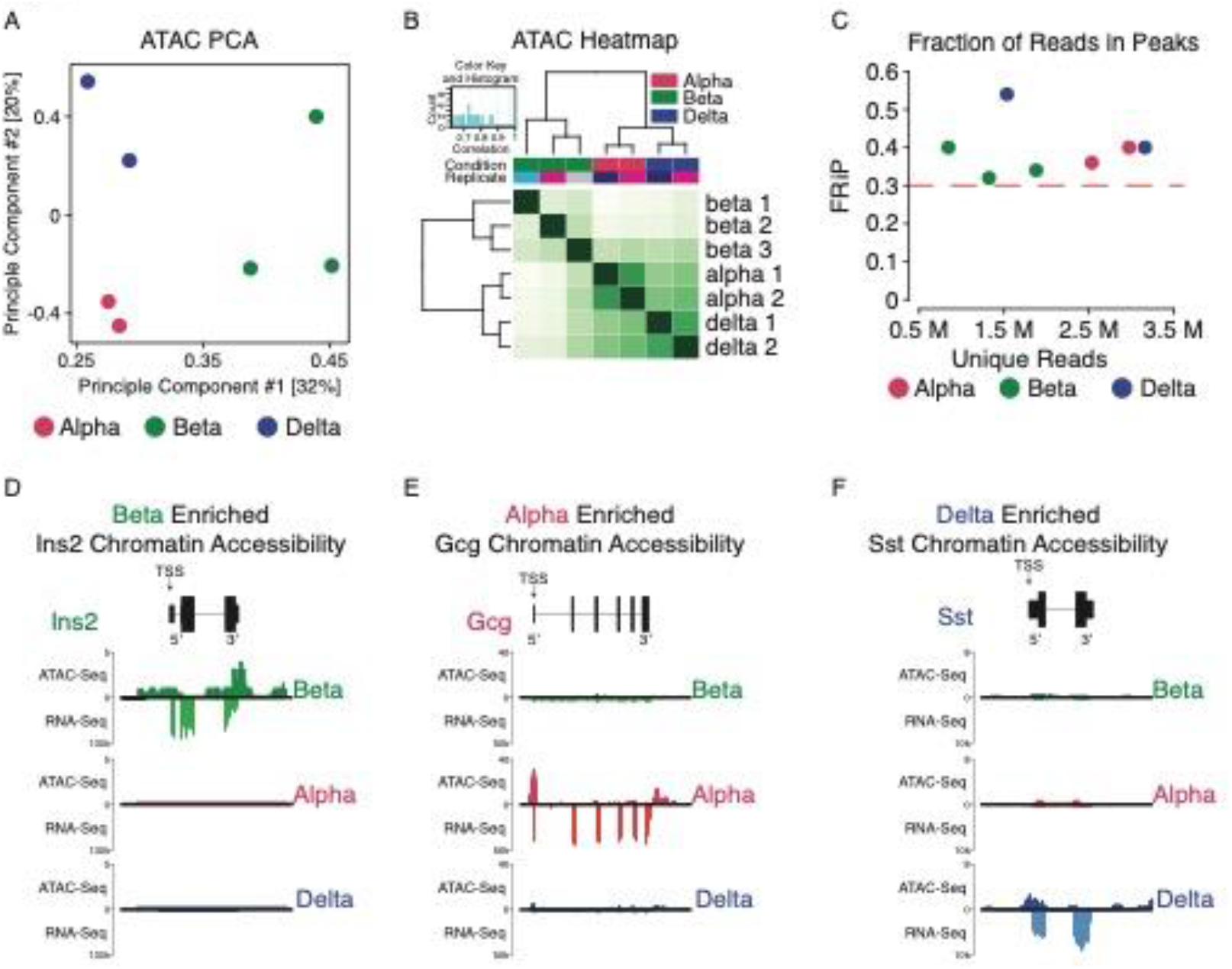
Validating alpha, beta, and delta chromatin accessibility ATAC Seq. A: Dimensional reduction through principal component analysis across seven samples from all three cell types (See Supplemental Table 1 for details). All three cell type’s replicates clustered closer together and separate from other cell types. B: Heatmap further confirming quality of replicates and similarity between replicates within each cell type. C: Fraction of Reads in Peaks (FRiP) score evaluation across samples, confirming high library complexity irrespective of depth of sequencing. D-F: Confirming chromatin accessibility at the TSS (arrows) against bulk RNA-Seq expression in key islet cell type-specific marker gene regions - *Ins2, Gcg,* and *Sst* - in beta, alpha, and delta cells, respectively.

### Downstream analysis

Transcription factor footprinting analysis and validation against existing ChIP data was performed through a modified script utilizing chromVar [59], regioneR [60], GenomicRanges [61], and motifmatchr [62] using the HOCOMOCO database [63]. Pathways analysis on differential chromatin accessibility was performed using the R Bioconductor packages ChIPseeker [64], ReactomePA [65], and clusterProfiler [66].

### Enhancer Identification

We developed a novel R package, *epiRomics*, to integrate our chromatin accessibility data alongside aggregated pancreatic islet ChIP and histone data to identify putative enhancer regions, as described [43]. The package, example data, and vignette can be found at: https://github.com/Huising-Lab/epiRomics and an interactive browser of the results from this manuscript is publicly available at: https://www.huisinglab.com/epiRomics_2021/index.html.

### Integrated data

Mouse alpha, beta, and delta (GEO: GSE80673), alongside alpha- and delta-transdifferentiated beta (GEO: GSE88778) transcriptomes were integrated into this analysis [6, 67]. Aggregated ChIP datasets of transcription factors and histone marks were added to the analysis through *epiRomics* [43] to identify putative enhancer regions (Supplemental Table 2).

## Results

### ATAC-Seq validation

To determine whether chromatin accessibility patterns differed between islet endocrine cell types, principal component analysis (PCA) was applied to peak calls across all samples. This confirmed that replicates clustered by cell type (Fig. 1A), a finding that was further validated by heatmaps using all defined peaks across replicates (Fig. 1B). Alongside quality control applied through the generation and analysis of this dataset, the fraction of reads in peaks (FRiP) score was in excess of the commonly applied benchmark of 30% (Fig. 1C). Furthermore, the FRiP score was independent of variability in unique read depth, indicating that peak calls were reproducible across all replicates within cell types independent of read depth range.

### Validation of islet cell chromatin accessibility data coupled to companion transcriptomes

After preliminary validation of our derived ATAC-Seq data, we checked for the presence of chromatin peak enrichment for alpha, beta, and delta marker genes that have been previously well-established and validated through complementary bench-lab or computational methods. We expected that if a gene is expressed within a cell type, its ATAC signal near the transcription start site (TSS) at promoter-proximal regions should reflect chromatin accessibility. Indeed, cell type-specific chromatin accessibility correlated with gene expression of *Ins2*, *Gcg*, and *Sst* genes for beta, alpha, and delta cells, respectively (Fig. 1D-F) [4, 6, 68]. After confirming chromatin accessibility in key cell-identity markers, we sought to compare and contrast select regions identified from prior groups that evaluated chromatin accessibility in human islets [5, 12, 38], as well as to further query whether chromatin was always uniquely enriched on a panel of cell type-specific genes across alpha (*Arx, Ttr, Gc)*, beta (*Ucn3, MafA, Pdx1)*, and delta cells (*Pdx1, Hhex, Rbp4, Ghsr*) (Fig. 2; Supplemental Fig. 2). Each of these genes demonstrated overall strong concordance between cell type-enriched gene expression and cell type-specific enrichment of available chromatin. This validated the utility of ATAC-Seq data to detect epigenetic factors that determine gene expression.

**Figure 2.**
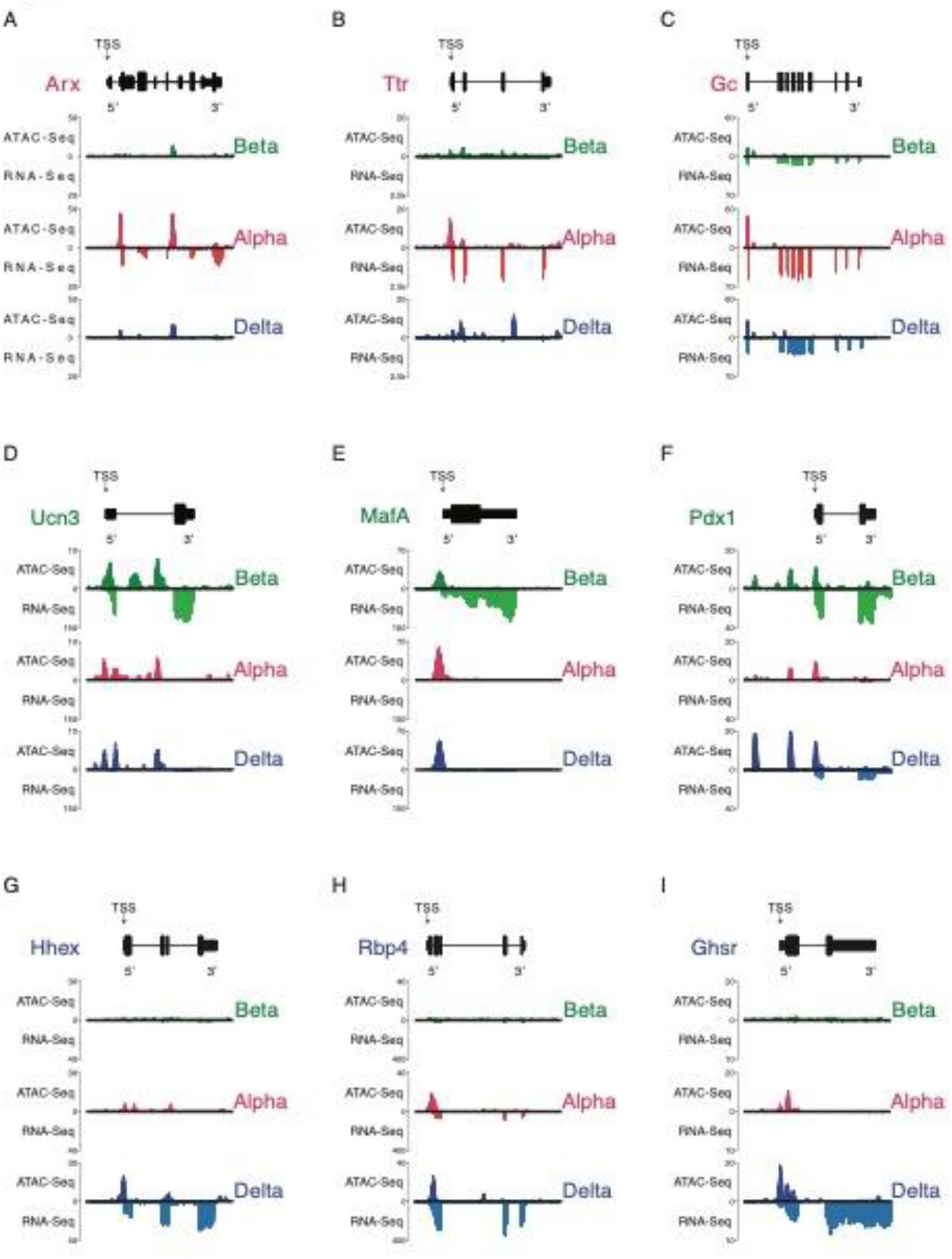
Validating chromatin accessibility ATAC Seq alongside companion RNA-Seq expression in alpha, beta, and delta cells against hallmark genes governing its respective cell’s identity. All genes are oriented for 5’ to 3’ end. A-C: Chromatin accessibility and transcript expression across alpha cell hallmark genes *Arx, Ttr,* and *Gc*. D-F: Chromatin accessibility and transcript expression across beta cell hallmark genes *Ucn3, Esr1,* and *Pdx1.* G-I: Chromatin accessibility and transcript expression across delta cell hallmark genes *Hhex, Rbp4,* and *Ghsr*.

### The chromatin landscape of the annotated genome across cell types

As genes make up a small fraction of the entire genome, we determined the overall distribution of peaks across the annotated genome within each cell type. We defined five regions of interest to further explore – promoter-proximal, intronic, exonic, downstream, or distal-intergenic (Fig. 3A). We identified a consensus set of 124,494 peaks marking open chromatin through the R package DiffBind. This number is comparable to the number of open regions found in previous studies of pancreatic islet chromatin accessibility [11, 38–40] (Supplemental Dataset 1). We then evaluated the distribution of called peaks present in at least one replicate within 3kb upstream of the TSS and confirmed that a majority of genes enriched in each islet cell type were accompanied by promoter-proximal peaks (Fig. 3B-D). The distribution of ATAC-Seq peaks across different pre-defined genomic areas was overall similar across alpha, beta, and delta cells. For each endocrine cell type between 21.98-24.88% of open chromatin was promoter-proximal, whereas promoter-proximal areas account for 2.41% of the mouse genome. A further 34.92-38.33% of peaks for all cell types were found on distal-intergenic regions, which was proportional to the fraction of the genome that falls into this category (Fig. 3E-G). Finally, we noted that between 33.07-33.65% of peaks occurred on intronic regions (first or other), relative to the 37.7% of the mouse genome classified as intronic [69].

**Figure 3.**
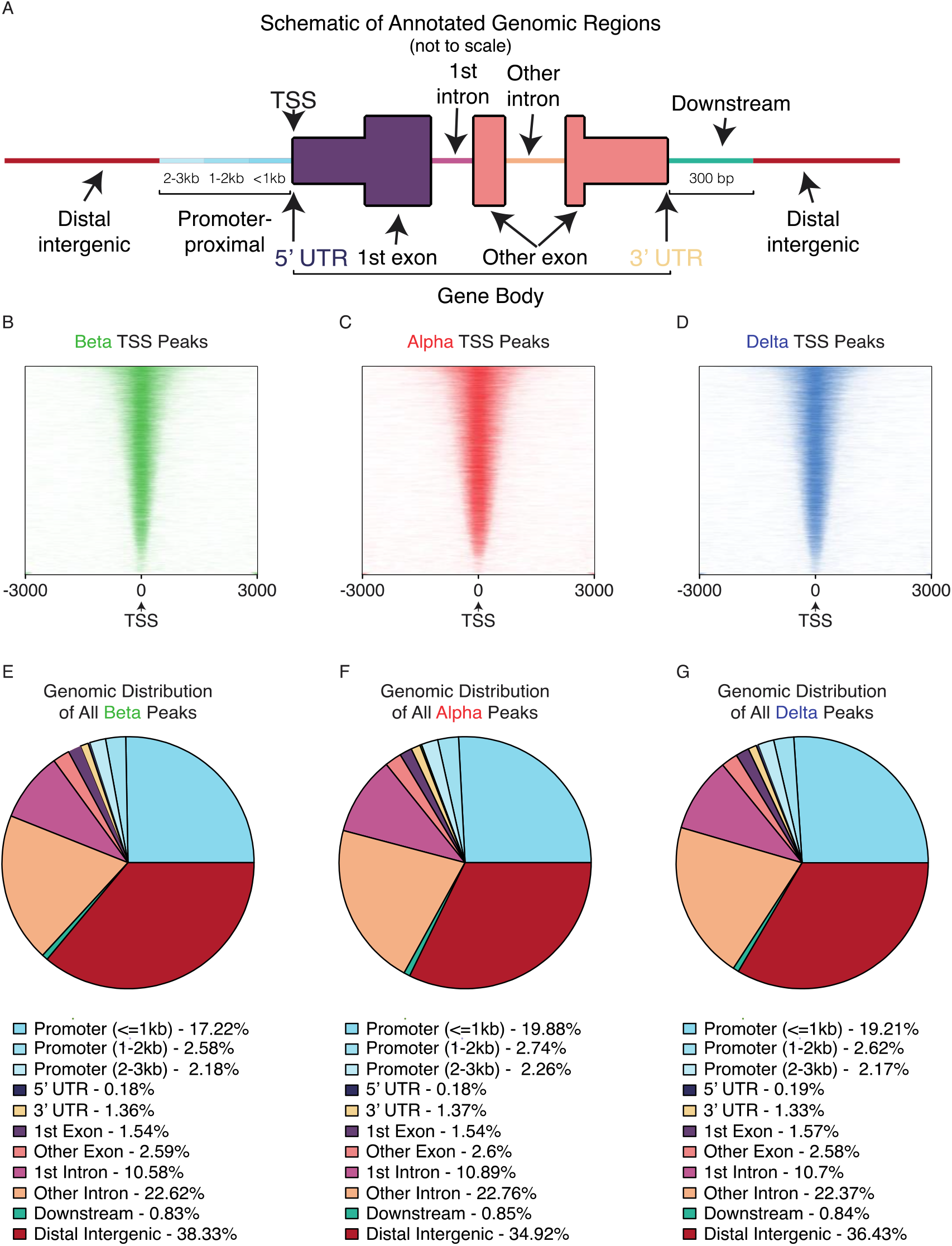
Evaluating chromatin accessibility ATAC Seq similarities and differences across all three cell types. A: Schematic of annotated genomic regions – promoter proximal, intronic, exonic, distal-intergenic, or downstream. B-D: TSS peak (defined as 3kb up or downstream each respective gene) chromatin accessibility density across beta, alpha, and delta cells. E-G: Distribution of chromatin peaks within each cell type across the annotated genome.

### Regional differences and characteristics of differentially enriched chromatin

As our overall distribution of ATAC-Seq peaks across different genomic regions was consistent across alpha, beta, and delta cells, we compared differential chromatin accessibility between these cell types in greater detail. To this end, we performed pairwise differential ATAC-Seq peak enrichment testing across alpha, beta, and delta cells. Out of 124,494 identified consensus regions of open chromatin across the three cell types, 18,409 (14.8%) differentially enriched peaks (p-value <= 0.05) were identified between alpha and beta (Fig. 4A), 12,722 (10.2%) between alpha and delta (Fig. 4B), and 16,913 (14.6%) between beta and delta cells (Fig. 4C).

**Figure 4.**
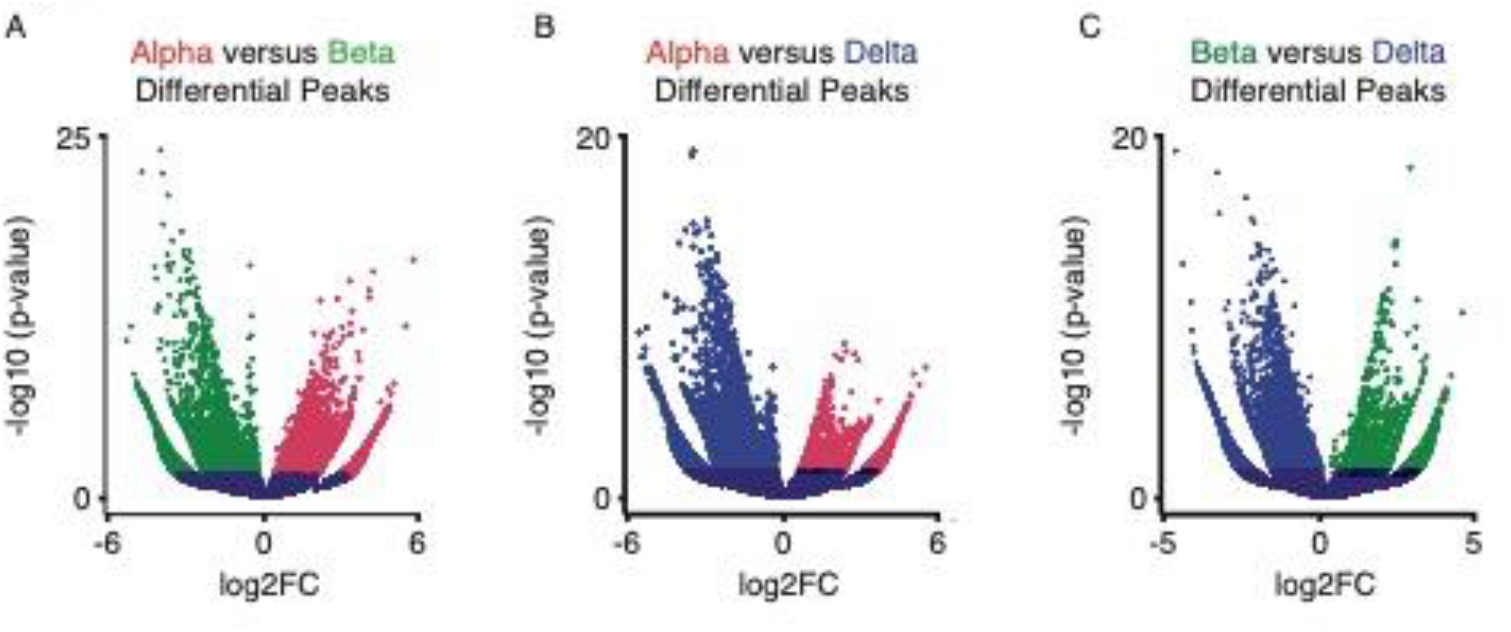
Comparing chromatin accessibility through differential enrichment analysis across alpha, beta, and delta cells. A: Differential chromatin accessibility peaks between alpha and beta ATAC Seq data. A total of 18,409 peaks were considered differentially enriched at p-value <= 0.05 (Supplemental Dataset 1). B: Differential chromatin accessibility peaks between alpha and delta ATAC Seq data. A total of 12,722 peaks were considered differentially enriched at p-value <= 0.05 (Supplemental Dataset 1). C: Differential chromatin accessibility peaks between beta and delta ATAC Seq data. A total of 16,913 were considered differentially enriched at p-value <= 0.05 (Supplemental Dataset 1).

After performing differential peak enrichment testing, we discovered that 22.89% of all differentially enriched peaks between alpha and beta cells were promoter-proximal (0-3kb) (Fig. 5A). A further 33.22% of differential peaks were linked to distal-intergenic regions and another 33.61% of differential peaks were intronic (first and other combined) (Fig. 5A). This assessment of differential peaks without considering the direction of enrichment revealed no major difference with overall peak distribution described earlier (Fig. 3). However, when factoring in the direction of enrichment we observed that 35.08% of alpha cell-enriched peaks was promoter-proximal. In contrast, only 12.5 % of beta cell enriched peaks occurred in promoter-proximal areas (Fig. 5B). Instead, a majority of ATAC peaks enriched in beta cells were located at distal-intergenic regions (45.41%) (Fig. 5B).

**Figure 5.**
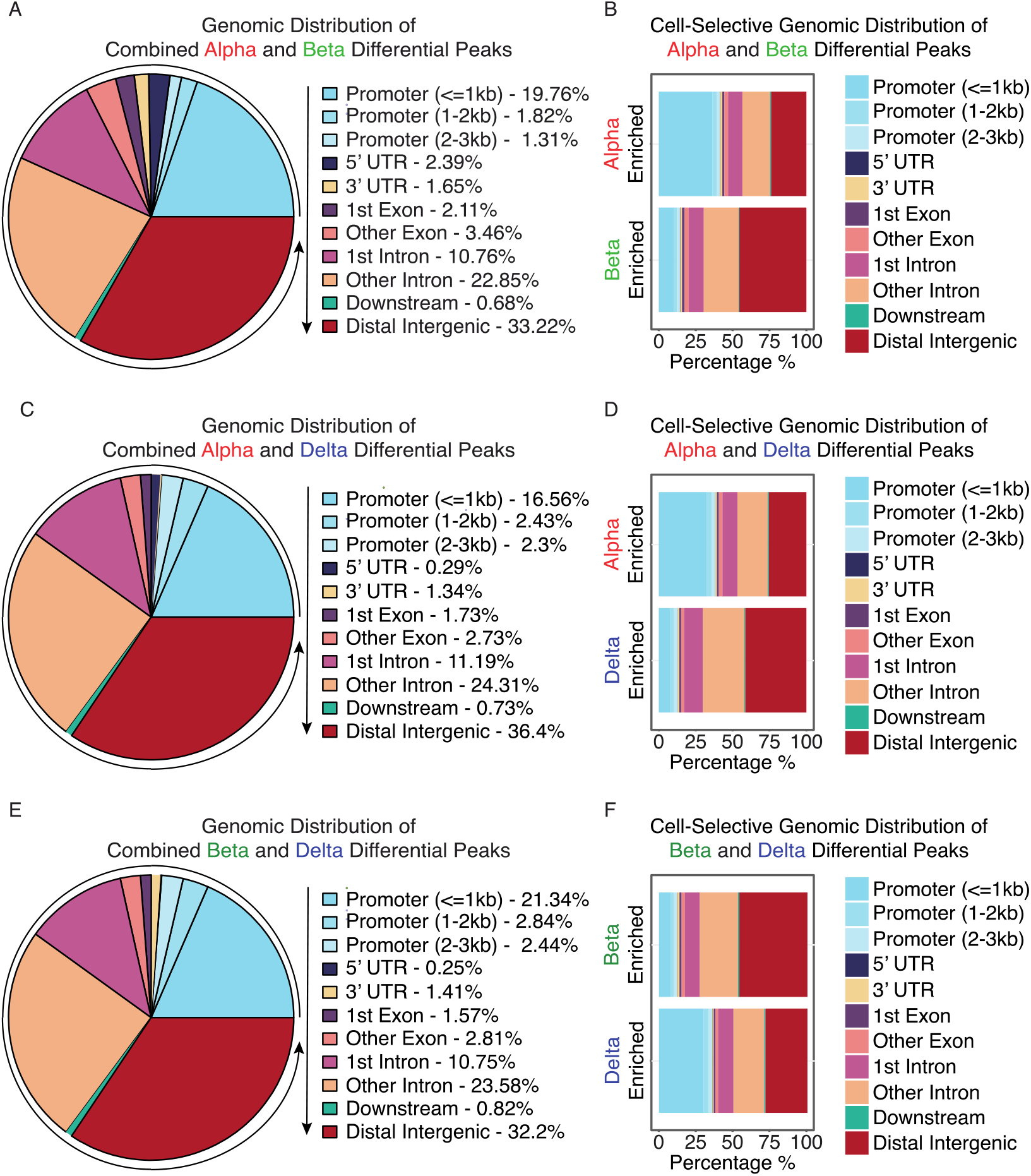
Regional differences and characteristics of differentially enriched peaks between alpha, beta, and delta cells. A: Distribution of regional preference across the annotated genome of differentially enriched peaks between alpha and beta cells. B: Regional preference breakdown of differentially enriched peaks between alpha and beta cells, indicating prevalence of enrichment for each cell type and genomic annotation. Differentially enriched chromatin favored promoter-proximal peaks in alpha cells, and distal-intergenic regions in beta cells. C: Distribution of regional preference across the annotated genome of differentially enriched peaks between alpha and delta cells. D: Regional preference breakdown of differentially enriched peaks between alpha and delta cells, indicating prevalence of enrichment for each cell type and genomic annotation. Differentially enriched chromatin favored promoter-proximal peaks in alpha cells, and distal-intergenic regions in delta cells. E: Distribution of regional preference across the annotated genome of differentially enriched peaks between beta and delta cells. F: Regional preference breakdown of differentially enriched peaks between beta and delta cells, indicating prevalence of enrichment for each cell type and genomic annotation. Differentially enriched chromatin favored promoter-proximal peaks in delta cells, and distal-intergenic regions in beta cells.

Between alpha and delta cells, we identified that 21.29% of differentially enriched peaks occurred promoter-proximally. Another 36.4% of peaks occurred on distal intergenic regions and 35.5% on intronic regions (Fig. 5C). A similar preference of alpha cell-enriched peaks in promotor-proximal regions was evident when comparing alpha to delta cells, with 30.33% of all enriched alpha peaks occurring promoter-proximally, but only 9.56% of delta cell peaks. Instead, 38.41% of delta cell enriched peaks were distal-intergenic (Fig. 5D).

Lastly, between beta and delta cells, 26.62% of all differentially enriched peaks were promoter-proximal, 32.2% distal intergenic, and 34.33% on intronic regions (Fig. 5E). Further break down revealed a bias towards distal-intergenic enriched peaks within beta cells (42.86%), as opposed to promoter-proximal peaks in delta cells (28.20%) (Fig. 5F).

### Differential chromatin enrichment in the majority of cases correlates with gene expression

So far, we detected a disproportionate fraction of peaks associated with promoter-proximal regions in general (Fig. 3). Moreover, ATAC-Seq peaks that were differentially enriched in alpha and - to a lesser extent - delta cells were considerably more likely to occur at promoter-proximal sites. Instead, peaks enriched in beta cells more likely occurred at distal intergenic regions (Fig. 5). Therefore, we determined whether the enrichment of promoter-proximal peaks correlated with increased expression of the corresponding. Genes with increased expression in a cell type accompanied by a significantly enriched ATAC-Seq peak proximal to its TSS were considered ‘congruent’ genes (Fig. 6A). The underlying mechanism in such a scenario might be the presence of transcriptional activators at the promoter-proximal site that promote gene expression. Conversely, genes with a significantly enriched ATAC-Seq peak proximal to its TSS accompanying a reduction in corresponding gene expression were considered ‘incongruent’ genes (Fig. 6B). The underlying mechanism for these genes might be the presence of transcriptional repressors at the promoter that prevent gene expression (Supplemental Dataset 2) [70–73]. Finally, genes that had significantly enriched chromatin in either cell type, but no evidence of mRNA expression were considered ‘unexpressed’ (Fig. 6C).

**Figure 6.**
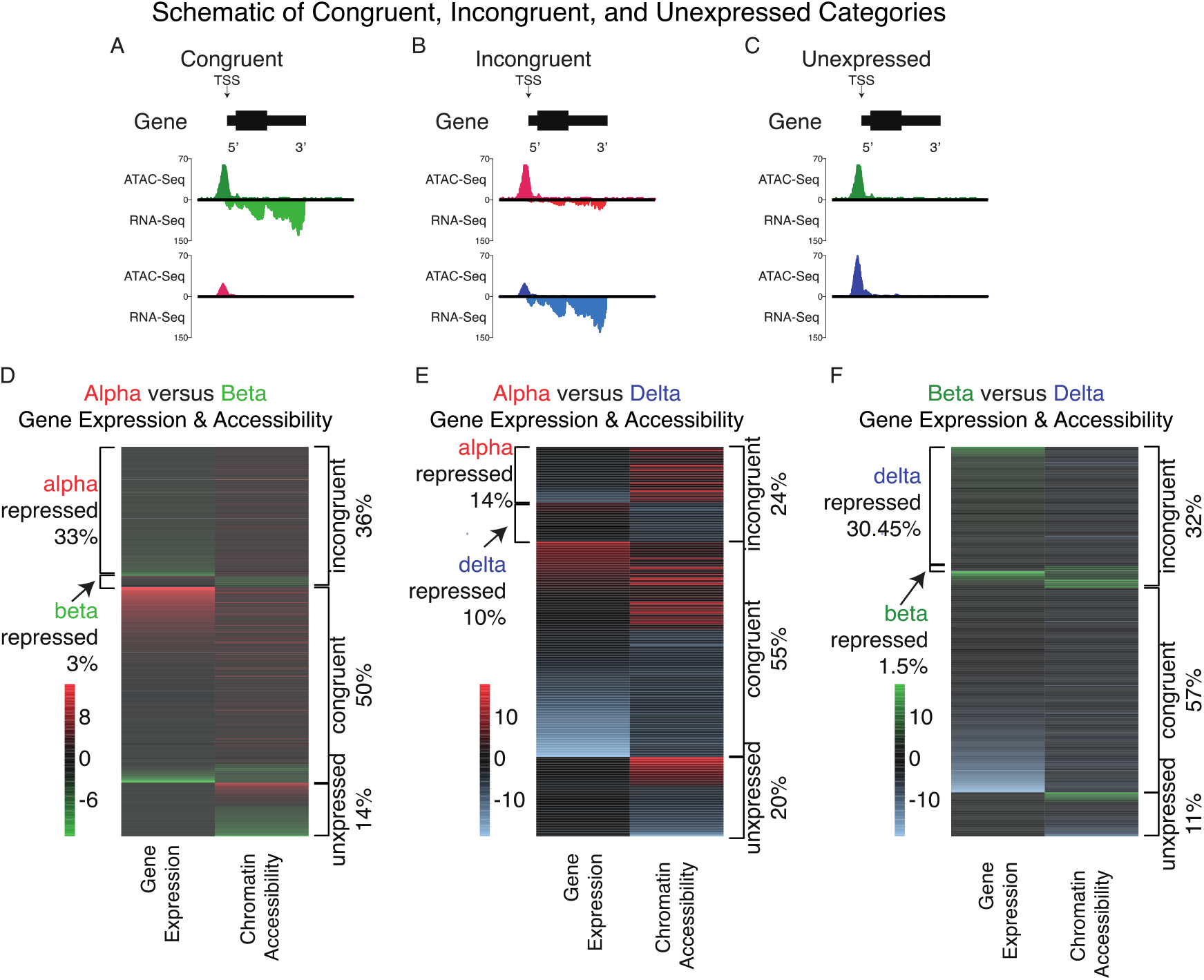
Differentially enriched chromatin at TSS genic regions and their respective gene’s expression between alpha, beta, and delta cells. A-C: Schematic of ‘congruent’, ‘incongruent’, and ‘unexpressed’ categories used to determine the association of enriched chromatin at TSS genic regions and respective gene expression. D: Differentially enriched chromatin at TSS genic regions and their respective gene’s expression between alpha and beta cells. The majority (50%) of genes with enriched chromatin at promoter-proximal regions around their TSS had correlated gene expression (congruent). Another 36% of chromatin enriched TSS regions showed repressed gene expression for each cell type (alpha repressed (33%) or beta repressed (3%)), and finally, 14% were unexpressed. E: Differentially enriched chromatin at TSS genic regions and their respective gene’s expression between alpha and delta cells. The majority (55%) of genes with enriched chromatin had correlated gene expression (congruent). Another 24% of chromatin enriched TSS genic regions showed repressed gene expression for each cell type (alpha repressed (14%) or delta repressed (10%)), and finally, 20% showed no expression. F: Differentially enriched chromatin at TSS genic regions and their respective gene’s expression between beta and delta cells. The majority (57%) of genes with enriched chromatin had correlated gene expression (congruent). Another 32% of chromatin enriched TSS regions showed repressed gene expression for each cell type (beta repressed (1.5%) or delta repressed (30.45%)), and finally, 11% showed no difference.

When we compared differentially enriched TSS-associated chromatin against corresponding gene expression between alpha and beta cells, we found that in the majority of cases (86%), differential chromatin enrichment on TSS regions successfully captured the epigenetics of gene regulation. Exactly, 50% of genes with differentially enriched chromatin at the TSS had a corresponding increase in gene expression within the same cell type (congruent genes). 36% showed TSS chromatin accessibility enrichment, but with a reduction in gene expression for each cell (incongruent genes - either alpha repressed (33%), or beta repressed (3%)). Strikingly, a substantial majority of the incongruent genes in this comparison were alpha repressed. Finally, only 14% of all genes with differentially enriched TSS chromatin showing no expression in either cell type (unexpressed) (Fig. 6D). A further visualization of select gene expression against TSS-associated chromatin accessibility indicated the majority as congruent, with a highlighted example of an incongruent (putatively alpha repressed or alpha cell poised) gene observed in the alpha cell TSS enrichment for the beta-specific genes *MafA* (Fig. 2E; Supplemental Fig. 3A). *MafA* is a key transcription factor enriched in beta cells yet shows abundant chromatin accessibility in alpha cells.

We observed a similar distribution between congruent (55%), incongruent (24%), and no expression genes (20%), between alpha and delta cells. We noted a more uniform distribution between alpha (14%) and delta (10%) repressed genes. (Fig. 6E). Upon visualizing gene expression and chromatin accessibility, we confirmed congruent gene expression and TSS chromatin accessibility of key transcription factors known to regulate both alpha – *MafB*, *Ttr*, and *Arx* – and delta – *Pdx1* and *Hhex* – cell fate (Supplemental Fig. 3B).

For our final pairwise comparison between beta and delta cells, we again found a similar fraction of congruent (57%), incongruent (32%), and no expression (11%) genes (Fig. 6F). We noted a minor fraction of repressed genes with open chromatin in beta cells (1.5%), with the overwhelming majority of repressed genes corresponding to delta cells (30.45%), similar to the pattern seen in alpha repressed genes between alpha and beta cells. Further visualization of select marker gene expression against chromatin accessibility showed generally good congruence between chromatin accessibility at the TSS and gene expression (Supplemental Fig. 3C).

### Poised genes are enriched in beta cells with a non-beta cell lineage history

To further interrogate whether these alpha- or delta-repressed genes could be poised beta cell genes, we incorporated transcriptome data from beta cells with an alpha- or delta-cell lineage history [67] – also from our companion RNA-Seq experiment. These cells, termed “transdifferentiated,” are functionally mature beta cells (defined by the presence of *Ucn3*), but have either a *Gcg-* or *Sst*-Cre lineage label, reflective of a lineage history as an alpha or delta cell, respectively. We reasoned that if alpha- or delta-repressed genes are poised beta cell genes, we should expect to observe a stepwise transition in gene expression levels, showing little or no expression in either alpha or delta cells, to intermediate expression in alpha- or delta-transdifferentiated cells, and full expression in beta cells. We confirmed that the majority (83.6%) of alpha-repressed genes showed intermediate expression in the alpha-to-beta-transdifferentiated population, and the highest expression in beta cells. A subset of genes (16.4%) showed the highest expression in the alpha-transdifferentiated population (Fig. 7A). We observed a similar pattern between delta, delta-transdifferentiated, and beta cells; however, only half (50.18%) of delta repressed genes demonstrated an intermediate expression in the delta-to-beta-transdifferentiated population and the highest in beta (Fig. 7B). The remainder of the genes showed the highest expression in delta-to-beta-transdifferentiated cells.

**Figure 7.**
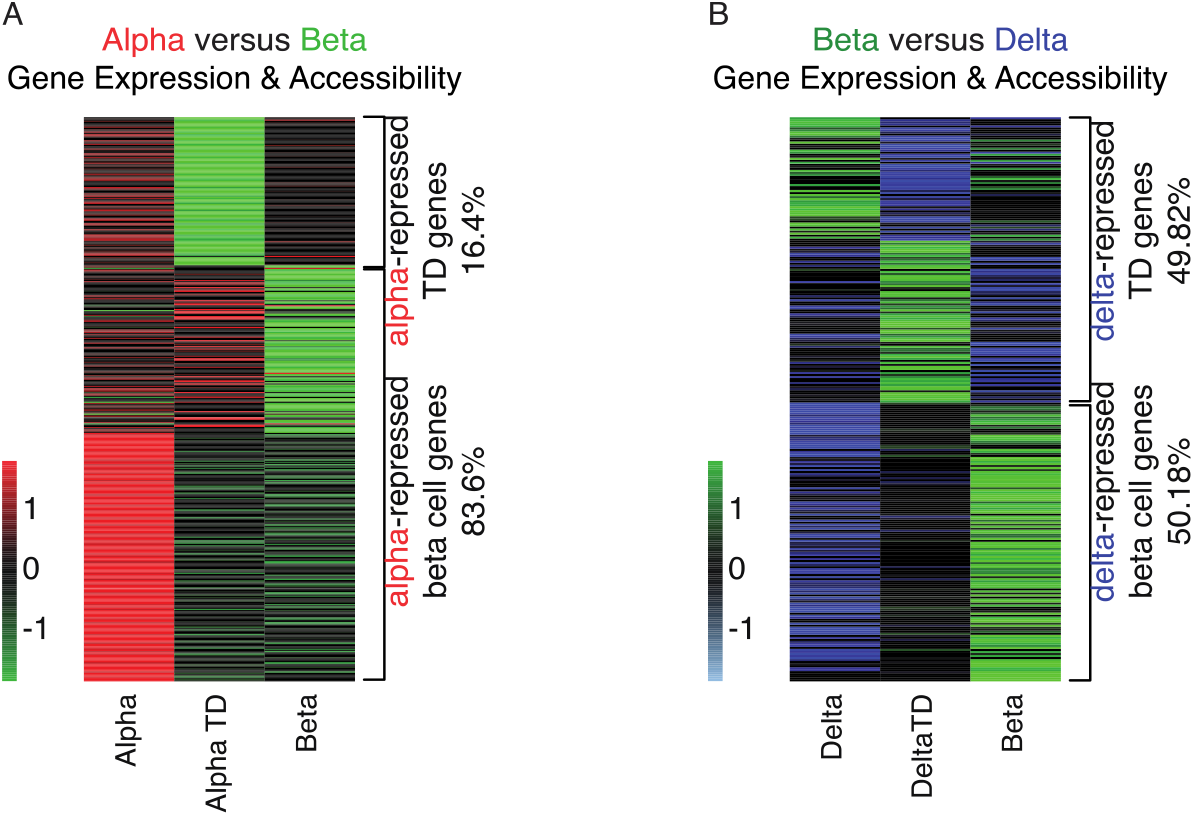
Gene expression of poised genes enriched in beta cells with a non-beta cell lineage. A: Evaluating alpha repressed genes (Fig. 6A) across alpha, alpha transdifferentiated, and beta cell transcriptomes. The great majority (83.6%) of genes repressed in alpha cells showed intermediate expression in alpha transdifferentiated cells, and highest expression in beta cells, further validating that alpha cells are poised to become beta cells, with a subset (16.4%) of those genes required for the transition. B: Evaluating delta repressed genes (Fig. 6C) across delta, delta transdifferentiated, and beta cell transcriptomes. Around half (50.18%) of genes repressed in delta cells showed intermediate expression in delta transdifferentiated cells, and highest expression in beta cells, validating that delta cells – to a lesser extent than alpha – are also poised to become beta cells, with the remainder of genes (49.82%) required for the transition.

### Differential meta-chromatin enrichment testing

Given that in a majority of cases, TSS-associated chromatin recapitulated the underlying regulation of gene expression, we inquired whether differentially enriched chromatin peaks were associated with genes concentrated in pathways or gene networks that would better reflect our understanding of the biology across these different islet endocrine cell types. Between alpha and beta cells, KEGG set pathway testing of differentially accessible chromatin identified pathways related to protein digestion and absorption and cell adhesion molecules unique to beta cells, Hippo, Wnt, and ubiquitin-mediated proteolysis unique to alpha cells, and MAPK, axon guidance, and cAMP pathways enriched within both (Supplemental Fig. 4A-B). Upon comparing the differentially accessibly chromatin between alpha and delta cells, adherens junctions appeared selective to delta cells, while no pathways were enriched specifically in alpha cells. MAPK, axon guidance, and Ras signaling pathways showed general enrichment of associated peaks within both alpha and delta cells (Supplemental Fig. 5A-B). Lastly, in beta and delta cells, a pairwise analysis of differentially accessible chromatin identified the Glycosaminoglycan (GAG) biosynthesis pathway as unique to beta cells - where GAG metabolism and biosynthesis impairment has been linked to beta cell dysregulation [74], adherens junctions and Rap1 signaling pathways unique to delta cells, and MAPK, axon guidance, and cAMP signaling pathways enriched within both (Supplemental Fig. 6A-B).

### Islet transcription factor ChIP-Seq binding correlates with open chromatin

After exploring the interrelationship between accessible chromatin and gene expression, we expanded our approach to include additional epigenetic controls to the regulation of islet cell gene expression. We therefore aggregated high-quality, mouse pancreatic islet transcription factor binding data via ChIP-Seq - Pdx1 [75], Nkx6-1 [76], Neurod1 [77], Insm1 [77], Foxa2 [77], Nkx2-2 [78], Rfx6 [79], MafA [24], Isl1 [80], Kat2b [81], Ldb1 [80], and Gata6 [82] - and asked what fraction of open chromatin – as defined by our consensus ATAC-Seq peak set – containing binding sites for each respective transcription factor. The transcription factors Foxa2 (29.07%), Insm1 (28.40%), and Neurod1 (20.09%) had the highest percentage of ChIP-confirmed binding site overlap with open chromatin. This provided further support that our ATAC-Seq data was of high quality and suggested that open chromatin is a reliable indicator of epigenetic regulation (Supplemental Table 3A). To further explore whether these aggregated transcription factor ChIP data convey epigenetic relevance, we queried what fraction of total ChIP binding sites overlapped with open chromatin. Indeed, we observed that in several cases, over 50% of transcription factor ChIP binding sites overlapped with open chromatin, with the transcription factors Nkx2.2 (63.79%), Neurod1 (55.07%), and Insm1 (51.49%) showing the greatest degree of overlap (Supplemental Table 3B). These results supported our findings that open chromatin reflects epigenetic regulation in alpha, beta, and delta cells, in part through the binding of transcription factors.

### Transcription factor motif finding suggests genomic preferences at differentially enriched chromatin between cell types

After observing a strong degree of overlap of known islet transcription factor binding on open chromatin, we conducted an unbiased evaluation whether DNA motifs for their respective transcription factor proteins were differentially enriched on ATAC peaks in promoter, intronic, exonic, downstream, or distal regions. We included transcription factors with known DNA-binding motifs to determine if they were more likely to occur at specific areas of the genome. We required that the transcription factor associated with the DNA sequence motif considered is expressed (RPKM>0) in the cell type with chromatin-motif association.

Motifs for key transcription factors involved in beta cell identity, such as MafA, were present ubiquitously across most functional regions we defined (promoter-proximal, intronic, exonic, downstream, and distal intergenic) (Fig. 8A). In contrast, the motifs for cell-identity drivers Irx2 [4] were concentrated at the promoter-proximal regions of chromatin peaks associated with genes differentially expressed by alpha cells. Insm1 [77] motifs were concentrated at the promoter-proximal regions of chromatin peaks associated with genes differentially expressed by beta cells. In another example, DNA-binding motifs associated with the ubiquitous islet transcription factor Pax6 [83] were concentrated on intronic chromatin.

**Figure 8.**
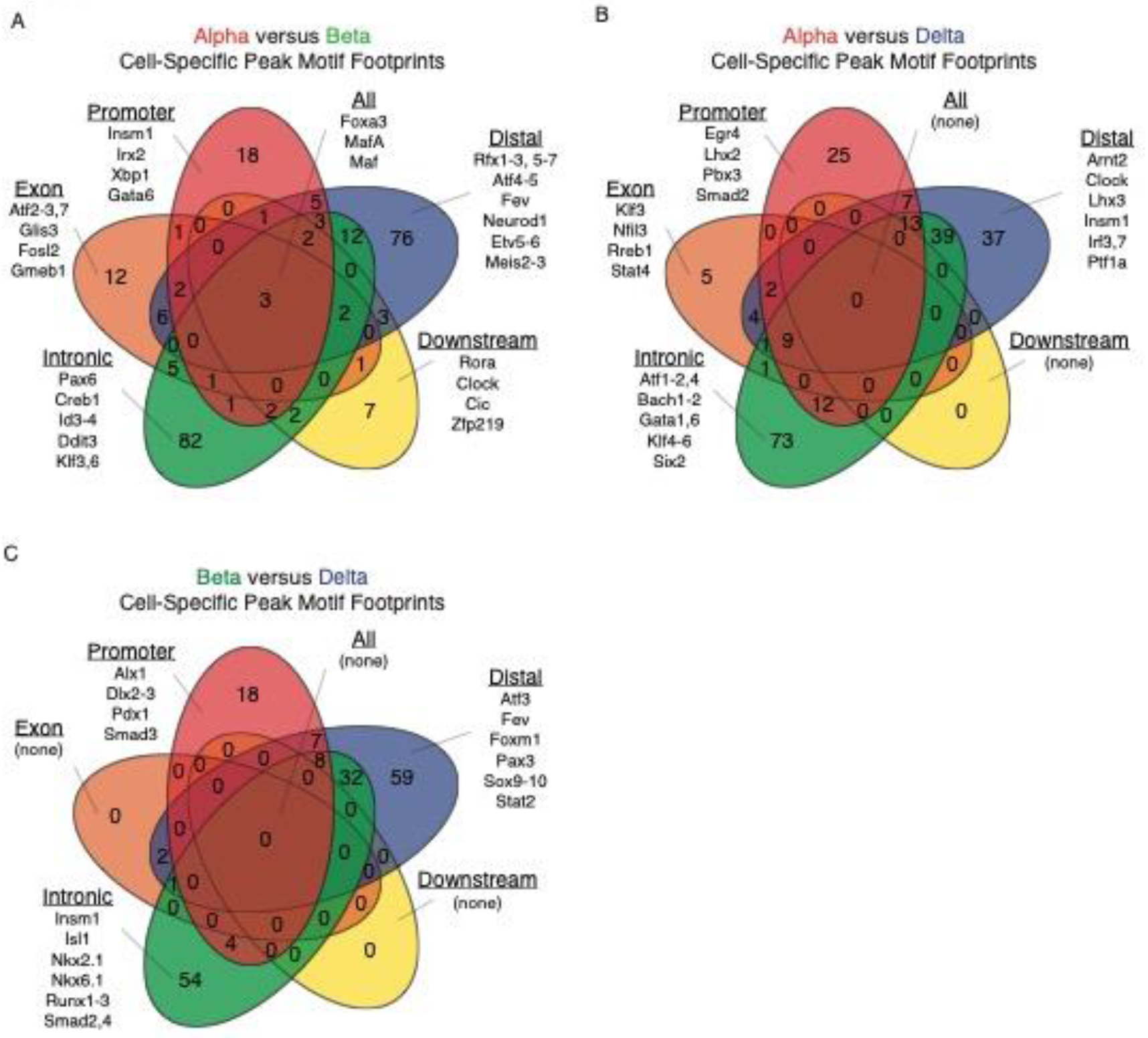
Evaluating expressed, cell-specific transcription factor footprints on differentially enriched peaks across cell types. A: Evaluating cell-specific transcription factor footprints on differentially enriched peaks for alpha and beta cells, suggesting transcription factor preference for these peaks across the functionally annotated genome. Notably, three known transcription factors were predicted to overlap all defined regions of the genome, whereas others showed preference for binding at either promoter, exon, intron, or distal regions, suggesting different mechanisms of regulation. B: Evaluating cell-specific transcription factor footprints on differentially enriched peaks for alpha and delta cells, suggesting transcription factor preference for these peaks across the functionally annotated genome. No known transcription factor was predicted to bind to all defined regions of the genome, with the majority binding to either intronic, distal, or promoter areas. C: Evaluating cell-specific transcription factor footprints on differentially enriched peaks for beta and delta cells, suggesting transcription factor preference for these peaks across the functionally annotated genome. No known transcription factor was found predicted to bind to all defined regions of the genome, with the great majority showing a preference for distal, intronic, or promoter regions.

We performed the same transcription factor footprinting test between alpha and delta cells (Fig. 8B). Of note, the motif for Pbx3, a transcription factor driving *Sst* expression in delta cells [84], was enriched in accessible chromatin at promoter-proximal regions. The motif for Stat4, recently implicated in establishing alpha cell identity [85] was concentrated exonic chromatin. Lastly, the motif for Ptf1a, a transcription factor identified in early pancreatic endocrine cell development [86], was preferentially associated with areas of open chromatin at distal intergenic regions.

Between beta and delta cells, no single transcription factor motif overlapped across all five functional categories, nor were there any unique to downstream or exonic regions, as we observed in the prior alpha and delta comparisons (Fig. 8C). Of note, the motif for Smad3, a transcription factor important for islet development [87], as well as the negative regulation of insulin secretion in beta cells via occupancy of the insulin promoter [88], was concentrated in promoter-proximal accessible chromatin. Motifs for Insm1 [77] and Nkx6.1 [89], both key beta cell identity transcription factors, were preferentially associated with accessible chromatin at intronic regions. Lastly, motifs for Fev – recently identified as important for the development and differentiation of the endocrine lineage [90] - and Atf3 *–* linked to enhancer regions in EndoC-bH1 cells [11] - were enriched in accessible chromatin at distal regions.

### Validating motif calls against aggregated islet ChIP datasets

As we observed motif binding site preferences across promoter, intronic, exonic, downstream, or distal chromatin regions, we wished to confirm how accurately predictive DNA motif binding sites conveys true transcription factor binding. To do so, we once again turned to our aggregated pancreatic islet ChIP datasets. We applied the same motif detection method as above on individual ChIP datasets and on all open chromatin – as derived from our ATAC-Seq consensus peak set –and assessed how well predicted motif binding overlapped with true ChIP peaks from our selected list of ChIP-Seq data. We observed strong (57%) true positive and low (8.34%) false negative values for the Rfx DNA-motif’s ability to predict all Rfx ChIP binding sites (Supplemental Table 4A). We then limited this same test to Rfx6 ChIP-Seq binding sites (35.65%) that were shared across 1.19% of all open chromatin ATAC peaks (Supplemental Table 3A-B). Notably, when comparing DNA-motif predictions for Rfx6 against these shared Rfx6 ChIP-Seq binding sites, we observed that 65.10% were true positives, while 8.81% were false predictions (Supplemental Table 4B). However, we also observed a broad distribution in true positive values across these DNA-binding motifs (4.71-65.10%), and also noted a relatively low range of false positives (1.68-8.81%). This is reflective of a limitation in the ability for DNA-motif’s to consistently predict true transcription factor binding.

### Determining overlap of differentially enriched chromatin with islet ChIP and histone datasets

Given that may of the DNA-motifs associated with transcription factors do not have available ChIP-Seq datasets derived from mouse pancreatic islets, we sought to understand whether differential chromatin between cell types could be associated to transcription factors and histone markers integral to pancreatic islet cell fate that do have available ChIP data, as opposed to relying only on predictive motifs. As we previously observed strong enrichment of transcription factor binding sites across all open islet chromatin, we wanted to confirm if this overlap is augmented in differentially enriched chromatin associated with pancreatic islet transcription factor binding sites via aggregated islet ChIP data - Pdx1, Nkx6-1, Neurod1, Insm1, Foxa2, Nkx2-2, Rfx6, and MafA - and select, key histone marks — H3K27ac [91], H3K4me3 [91], and H3K4me1 [12]. Our intention in integrating these data was in anticipation that they may help delineate whether enhancer regions are poised (defined by: H3k4me1) [92], or active (defined by: H3k27ac and H3k4me1) [93], and whether promoter regions are active (defined by: H3k4me3) [94]. Indeed, we observed that differentially enriched peaks within our comparisons occurred at much higher rates than random chance across a majority of transcription factor ChIP data associated with islet cell identity (Supplemental Fig. 7A-C) as well as all predictive histone marker regions (Supplemental Fig. 7D-F). This further supported our hypothesis that open chromatin, and now specifically differentially enriched chromatin, would be directly associated with the transcription factors responsible for shaping islet cell-specific gene expression patterns and identity. As differentially enriched chromatin is associated with cell-identity regulatory networks, we inquired to selectively evaluate these regions for enhancers that may be relevant to pancreatic islet cell identity.

### Identifying and visualizing putative islet cell-type specific enhancers via epiRomics

These transcription factor and histone ChIP datasets were then fed into an R package named *epiRomics* that we developed to identify putative enhancer regions involved in pancreatic islet cell identity. We defined enhancer regions by the co-localization of H3k27ac and H3k4me1 histone modifications within islet cell chromatin. We stringently narrowed our definition further by requiring these regions to also have transcription factor binding sites - Pdx1, Nkx6-1, Neurod1, Insm1, Foxa2, Nkx2-2, Rfx6, MafA, Isl1, Kat2b, Ldb1, and Gata6 – defined by islet ChIP data that are either ubiquitously or selectively expressed across the three islet cell types (Fig. 2E-F; Supplemental Fig. 2E; Supplemental Fig. 8A-I). This first pass resulted in 28,647 putative enhancer regions (Supplemental Dataset 3). We then filtered this list against chromatin accessible regions from our ATAC-Seq data sets of alpha, beta, and delta cells, resulting in 16,651 putative active enhancers (Fig. 9A). To further increase our confidence in these enhancer calls, we crossed our putative enhancer regions against the curated FANTOM5 curated enhancer database [95]. This resulted in a conservative list of 3,535 putative enhancer regions. Of these 2,347 were inaccessible to at least two out of three islet endocrine cell types (Fig. 9B) (Supplemental Dataset 4).

**Figure 9.**
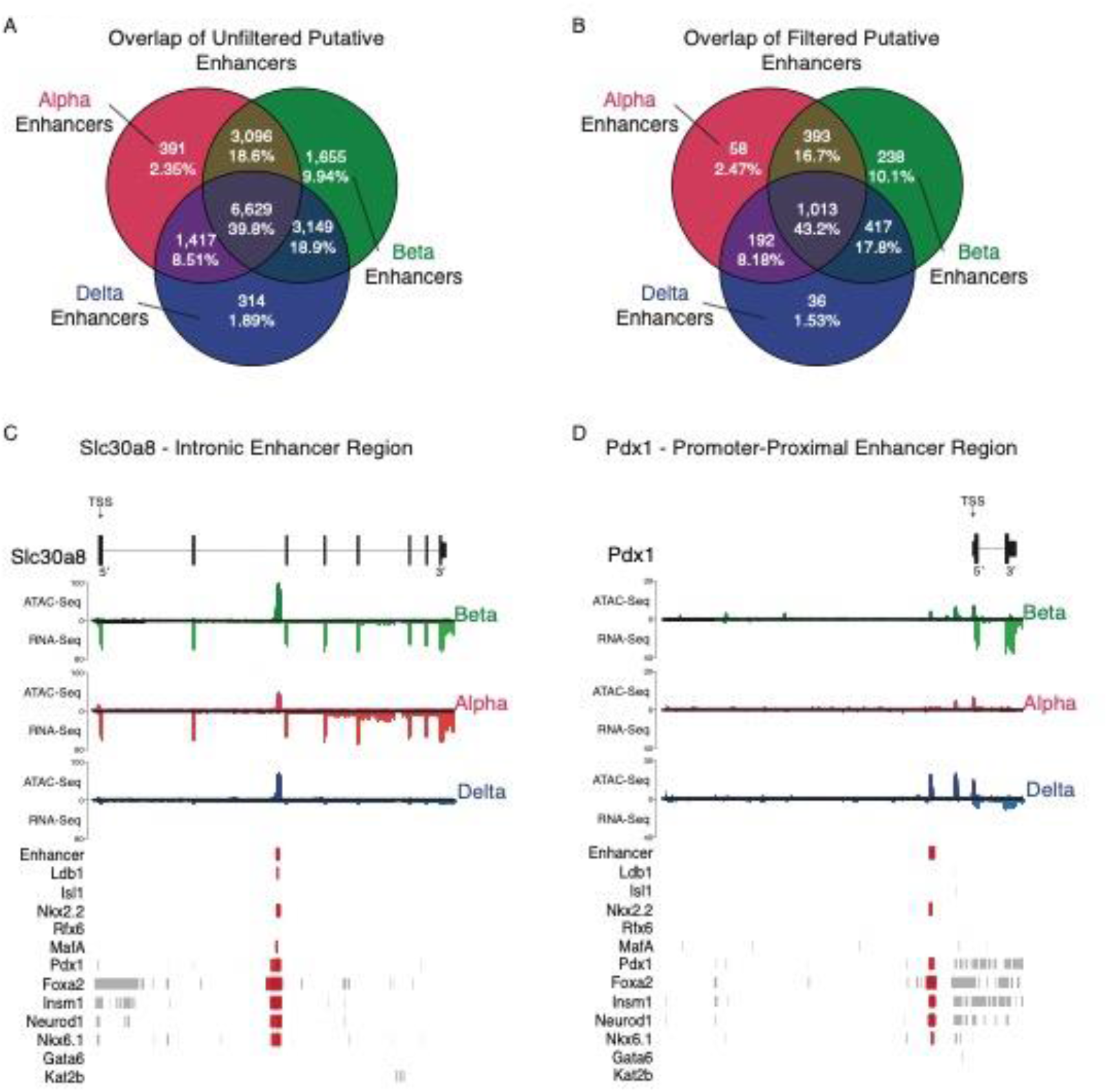
Putative enhancer detection overlap between the three cell types. A: First-pass overlap of unfiltered putative enhancers called with our novel package, epiRomics. Open chromatin regions in at least one cell type were crossed against two informative histone marks - H3k27ac and H3k4me1 – and transcription factor binding data to call putative enhancer regions. A total of 28,647 regions were identified (Supplemental Dataset 3). 39.8% of putative enhancer calls had chromatin accessible to all three cell types, suggestive of pancreatic endocrine cell development and maintenance involvement. The overlap of enhancer calls with open chromatin between any two cells type was 8.51% - 18.9%. Between 1.89% - 9.94% of calls were unique to one cell type alone. B: First-pass enhancer calls were filtered against the curated FANTOM5 database delineating all identified enhancers in the mouse genome. This resulted in a much more conservative list of 3,535 regions identified (Supplemental Dataset 4). The distribution of enhancers unique or common between cell types remained comparable, with 43.2% identified across all three cell types, and 1.53% - 10.1% unique to a cell type. C: Confirming an enhancer on the second intron of *Slc30a8*, identified in a previous study, with 14 sites of co-binding from multiple transcription factors. D: Confirming an a promoter-proximal enhancer (∼1kb upstream) of the gene that codes for the transcription actor *Pdx1*, with 9 sites of co-binding from multiple transcription factors.

In both putative enhancer lists, we found that 39.8-43.2% of the enhancer regions we identified were common across all cell types, supporting the theory that related cell types of a common origin would have a sizeable commonality of similar regulatory regions involved in development and maintenance (Fig. 9A-B). Interestingly, between 1.53–10.1% of called enhancers were associated with accessible chromatin unique to each cell type. Enhancer regions selective to beta cells were identified at the highest frequency (∼10%), while alpha and delta enhancers made up ∼2% of the list.

Upon evaluating whether or not our putative enhancer list would recapitulate two previous mouse pancreatic islet studies delineating enhancers, we confirmed that our approach was able to independently identify an established intronic enhancer on the *Slc30a8* gene, demonstrated to be regulated in part by the Pdx1 transcription factor (Fig. 9C) [96]. Our approach also supported a previously identified promoter-proximal enhancer region targeting *Pdx1*, with co-occurring binding sites for islet transcription factors Insm1, Neurod1, and Foxa2 (Fig. 9D) [77].

Given that our approach corroborated enhancers identified through complementary methods in previous mouse islet studies, we investigated cell-specific or common putative enhancer candidates by evaluating those with the highest number of transcription factor co-binding sites from our list. One of the top predicted beta cell-unique putative enhancer regions is located on the sixth exon of the *Slc35d2* gene and aligns with eight different ChIP co-localization binding sites (Fig. 10A). An alpha cell-unique putative enhancer located at a distal-intergenic region ∼30kb upstream of *Dusp10*, overlapped precisely with six sites of co-binding from various transcription factors (Fig. 10B). A delta cell-unique region at a distal-intergenic enhancer region ∼21kb upstream of *Gm20745* aligned closely with no fewer than 12 sites of co-binding from multiple transcription factors. (Fig. 10C). And finally, a common enhancer located ∼32kb upstream of *Snap25* – a gene expressed in alpha, beta, and delta cells, associated with a total of 17 co-binding sites of aggregated transcription factors. (Fig. 10D).

**Figure 10.**
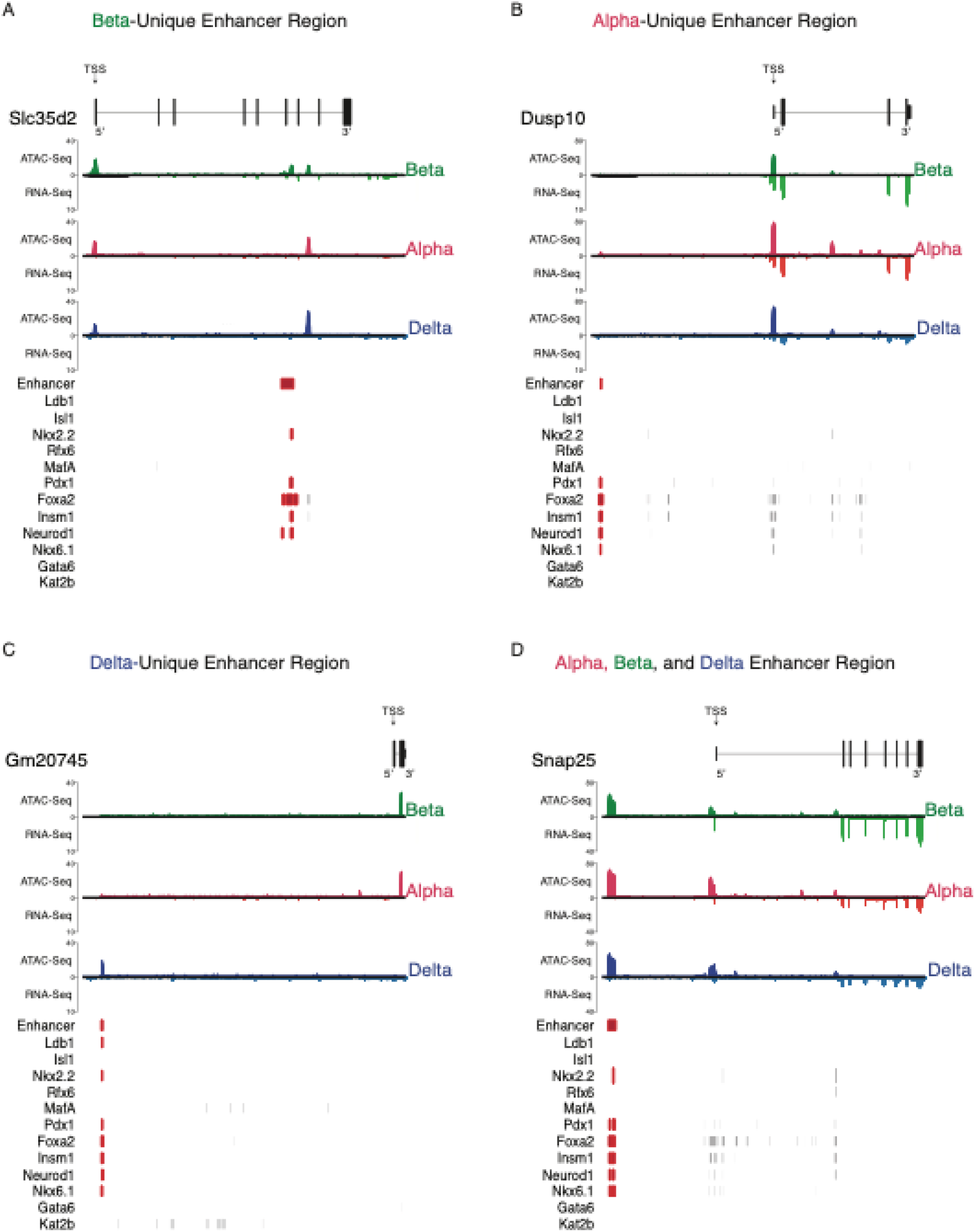
Visualizing novel, putative enhancer detection between cell types. A: Visualizing a beta- unique enhancer region. An exonic enhancer region selected from our filtered enhancer call list, with 8 sites of co-binding from various transcription factors relevant to pancreatic islet cell identity and maintenance [1]. B: Visualizing an alpha-unique enhancer region; a distal-intergenic enhancer region (∼30kb upstream of *Dusp10*) selected from our filtered enhancer call list, with 6 sites of co-binding from various transcription factors. C: Visualizing a delta-unique enhancer region. A distal-intergenic enhancer region (∼21kb upstream of *Gm20745*) selected from our filtered enhancer call list, with 12 sites of co-binding from various transcription factors. D: Visualizing a non-unique enhancer region common across all three cell types. A distal-intergenic enhancer region (∼32kb upstream of *Snap25*) selected from our filtered enhancer call list, with 17 sites of co-binding from various transcription factors.

We noted further examples of enhancer regions that are inaccessible to beta, but present in both alpha and delta (Supplemental Fig. 9A) cells, or others with chromatin accessibility across all cell types with an adjacent, intronic enhancer region uniquely available to beta cells alone (Supplemental Fig. 9B). In particular, the *Slc2a2* gene shares common open chromatin across alpha, beta, and delta cells. However, beta cells have a gained accessible chromatin region on the first intron identified as a putative enhancer and which overlaps with six co-binding sites. Finally, we noted more putative regions that were enriched in both alpha and beta cells, and present in delta cells as well (Supplemental Fig. 9C-D).

## Discussion

The high quality ATAC-Seq data derived from this study is the first dataset of its kind from FACS-purified mouse alpha, beta, and delta cells. Moreover, while bulk-ATAC-Seq data from human alpha and beta cells have previously been reported, our data are the first to report on chromatin accessibility of delta cells and combine these cell’s chromatin landscapes to companion transcriptome data. We believe that our data will provide a useful resource that complements our companion transcriptome data that we reported previously using the exact same combination of reporter strains [6]. Leveraging these data allowed us to confirm previous findings in a human ATAC-Seq study evaluating alpha and beta cells, which suggested that alpha cells are poised but repressed from becoming beta cells [5], and present evidence that supports that delta cells might be similarly epigenetically poised to adopt a beta cell like gene expression pattern. We also now harmonized our ATAC-Seq and RNA-Seq data with a wealth of -omics levels data from our colleagues, resulting in a comprehensive multi-layered omics overview that includes histone modifications and transcription factor binding sites. Finally, we made these data accessible through an intuitive interface that we developed to be navigated without any bioinformatics experience.

In evaluating the chromatin landscape of alpha, beta, and delta cells, we noted that over half of accessible chromatin in any of the cell types corresponded to promoter-proximal regions (∼25%) and intronic regions (∼32%), even though a much smaller fraction of the genome is represented by promoter-proximal sites. This underscores that a substantial portion of regulatory activity occurs directly at genic regions themselves. The enrichment of promoter-proximal and intronic open chromatin we observed in mouse islet cells agrees with previous findings in human studies [5, 38]. The strong presence of intronic peaks supported previously established findings of how enhancers on introns can act as suppressors [33] or drivers of gene expression [97] – in one instance, how *Pdx1* regulates the expression *Slc30a8* through an intronic enhancer [96] – and suggested that these intron regions of accessible chromatin may play a role in cell identity (Fig. 8D). Our findings of a large number of peaks residing at distal-intergenic regions (∼35%) agree with previous research identifying and emphasizing the role of distal intergenic regions acting as enhancers (Fig. 10C-D, Supplemental Fig. 9C-D) in pancreatic islet identity and functional beta cell behavior, and through linkage with T2D GWAS studies that link these regions to beta cell dysregulation [19, 20, 23, 24, 98, 99].

Upon evaluating differences in chromatin accessibility between pairwise comparisons, we discovered that overall, differentially enriched ATAC-Seq peaks in alpha or delta cells were more likely to occur at promoter-proximal regions adjacent to the TSS, whereas peaks enriched in beta cells were often found in distal intergenic or intronic regions, suggesting different mechanisms regulating alpha and delta cell fate specification (Fig. 5B, 5F). When comparing differentially enriched TSS-associated chromatin and respective gene expression, we observed a strong association between chromatin accessibility and gene expression. However, both alpha and delta cells showed a preference in putatively poised genes when either was compared to beta cells. Of note, *MafA* is a key transcription factor enriched in beta cells that shows abundant chromatin accessibility in both alpha and delta cells (Fig. 2E) but is only expressed in beta cells. Another notable example is *Pdx1*, which shows poised TSS enrichment in alpha cells, but is only expressed in beta and delta cells (Fig. 2F).

Moreover, a majority of alpha- and delta-repressed genes showed intermediate expression in the transdifferentiated populations (Fig. 7A-B), further supporting that these are indeed putatively poised. These observations are in line with prior data that suggest that alpha cells are epigenetically poised to become beta cells, but are prevented from assuming beta cell transcriptional programs by repressive regulators at key beta-specific transcription factors [5, 15]. Our observations here also fit reports of adult or juvenile transdifferentiation of alpha-to-beta, or delta-to-beta, respectively [5, 67, 100], although the contribution of these processes to beta cell regeneration is uncertain [101].

After evaluated motif binding on differentially enriched chromatin, we found that Irx2 and Insm1 motifs are enriched at promoter-proximal regions of cell-specific alpha peaks when comparing alpha and beta cells, suggesting that they directly drive gene expression or repression in alpha cells by binding to uniquely accessible chromatin [4, 77]. For Irx2, this indicates that it directly drives gene expression or repression in alpha cells. For Insm1, which is expressed more uniformly in all three endocrine cell types, the role that it plays at promoter-proximal accessible chromatin is more complex and cannot be as readily inferred. Between alpha and delta cells, the DNA binding motif for Pbx3, a transcription factor implicated in driving *Sst* expression in delta cells, was found preferentially enriched in accessible chromatin associated with promoter-proximal peaks [84], while the DNA motif for *Atf2*, identified as an enriched alpha cell motif in a previous human study, was found to be preferential enriched to chromatin associated with intronic peaks [5]. Between beta and delta cells, Insm1 and Nkx6.1 had motifs enriched at intronic chromatin regions [77, 89], while Fev – recently identified in pancreatic islet development - and Atf3 *–* linked to enhancer regions in EndoC-bH1 cells - were identified as preferential to chromatin associated with distal-intergenic regions via their DNA binding motifs [11, 90].

Utilizing the R package *epiRomics*, we were able to derive a set of 16,651 putative enhancer peaks. Of these regions, 16.7% of enhancers were shared between beta and alpha cells, and 17.8% were shared between beta and delta cells, as opposed to the 8.18% shared between alpha and delta cells. Of note, our approach identified previously identified intronic enhancers, such as the one located on *Slc30a8 that is* regulated in part via the binding of the transcription actor Pdx1 and a promoter-proximal enhancer region upstream of the Pdx1 that is associated with Insm1, Neurod1, and Foxa2 binding, also identified by our approach (Fig. 9D) [77]. One final example of an enhancer is situated on the first intron of *Slc2a2.* having unique chromatin accessibility to beta cells, coupled with multiple transcription factor binding sites, including Pdx1, MafA, and Nkx6.1, could possibly explain the expression of the gene in beta cells while it is near undetectable between alpha and delta cells (Supplemental Fig.9B). *Slc2a2* plays a necessary role glucose-stimulated insulin secretion [102], with a recent study identifying a downstream enhancer regulating *Slc2a2* requiring the co-occupancy of both MafA and Neurod1, but also noting that complex epigenetic interactions occur beyond the scope of this distal region [103].

One limitation of our approach was that we were constrained to using protein data available to the field. The substantial majority are transcription factors associated with beta cells, with the results reflective of this limitation. For instance, 10.1% of the enhancer regions called were unique to beta cells, whereas we were only able to identify 1.53-2.47% unique to either delta or alpha cells (Fig. 2E-F; Supplemental Fig. 2E; Supplemental Fig. 8A-I). The over-representation of beta cell-specific enhancer regions is probably explained by the fact that ChIP data for alpha and delta cell-specific enhancers obtained from pure populations of primary alpha and delta cells does not exist. While the majority of the transcription factors here are associated with beta cells, these data are still informative as delta and alpha regions with absence of beta-cell transcription factor binding may be areas regulated through other layers of epigenetics, such as methylation, or via alpha- or delta-specific transcription factors for which no ChIP data is currently available [94]. While further validation of these regions lays beyond the scope of this study, such information would be readily integrated in the future in the multi-omics resource we described here.

In conclusion, we provide a comprehensive snapshot of the characterization of chromatin similarities and differences between mouse alpha, beta, and delta cells. Here, we identify certain TSS genic regions that present as putatively poised in either alpha or delta cells and demonstrate intermediate expression of these genes in beta cells of a non beta-lineage (either alpha or delta transdifferentiated). We also provide a novel approach to identify active enhancers in these cell types through the use of these data alongside data integrated from the field using our package, *epiRomics*, first confirming enhancers identified in previous studies, and then showcasing novel regions with potential for further exploration. Taken together, we have demonstrated that the integration of chromatin accessibility data via ATAC-Seq with other epigenomic data can help further delineate regulatory regions and help answer outstanding questions in the field. Studies and resources such as these are relevant in such that they also function as a supportive resource for integrative research. Given this, we have made these data along with those aggregated through our approach as an interactive resource available on our website.

## Conclusion

Here we have established a comprehensive picture of chromatin accessibility between major islet endocrine cell types and present the novel chromatin landscape of delta cells. We identified differential chromatin accessibility at promoter-proximal regions in both alpha cells and delta cells, when compared to beta cells. This finding was in line with a previous study in human islets, and further builds on previous literature in the field suggesting that both alpha and delta cells can transdifferentiate into beta cells. We also identified preferentially binding pattern differences across the annotated genome in transcription factor DNA-motifs across differentially enriched chromatin. Our evaluation of whether chromatin enrichment at the gene body is always correlated with gene expression enrichment also demonstrated that transcriptional regulation plays a role in determining cell fate rather than chromatin dynamics alone. Lastly, we devised and provided a simple approach to utilize and integrate a subset of these epigenomic datasets – ChIP and histone - alongside our ATAC-Seq chromatin data integrated with our previously published transcriptome of the FACS-purified alpha, beta, and delta cells through the development of an R package, *epiRomics*. This allowed for the visualization of integrated epigenomic data, and furthermore applies a novel approach to identify putative enhancer regions, enabling a high-resolution overview of key regions that may be responsible for driving cell fate decisions in pancreatic islet cell types. We have made this an interactive resource publicly available at https://www.huisinglab.com/epiRomics_2021/index.html. We believe our data and the tool we developed to visualize it to be a valuable resource to our field in pursuit of a full understanding of the epigenetic control over islet gene expression.

## Acknowledgements

Conceptualization, Methodology, Investigation and Review, A.M.M., M.O.H, T.v.d.M.; Experimental Design, A.M.M., T.v.d.M., M.O.H.; Writing and Editing, and Visualization, A.M.M. and M.O.H.; Formal Analysis, A.M.M.; Supervision, Project Administration, Mentorship, and Funding Acquisition, M.O.H. M.O.H. received funding and consulting fees from Crinetics Inc. The other authors report no conflicts of interest. M.O.H is the guarantor of this work and takes responsibility for the integrity and accuracy of the data analyzed within this manuscript.

This work was supported by grants from the National Institutes of Health (NIDDK110276) and the Juvenile Diabetes Research Foundation (CDA-2-2013-54 and 2-SRA-2021-1054-M-N) to M.O.H. A.M.M. was supported by the Stephen F. and Bettina A. Sims Immunology Fellowship and the AWS Machine Learning Research Award. This work was also supported by the University of California, Davis Flow Cytometry Shared Resource Laboratory with funding from the NCI (P30 CA093373) and NIH S10 (OD018223) awards.

## Supplemental Tables

**Supplemental Table 1.**
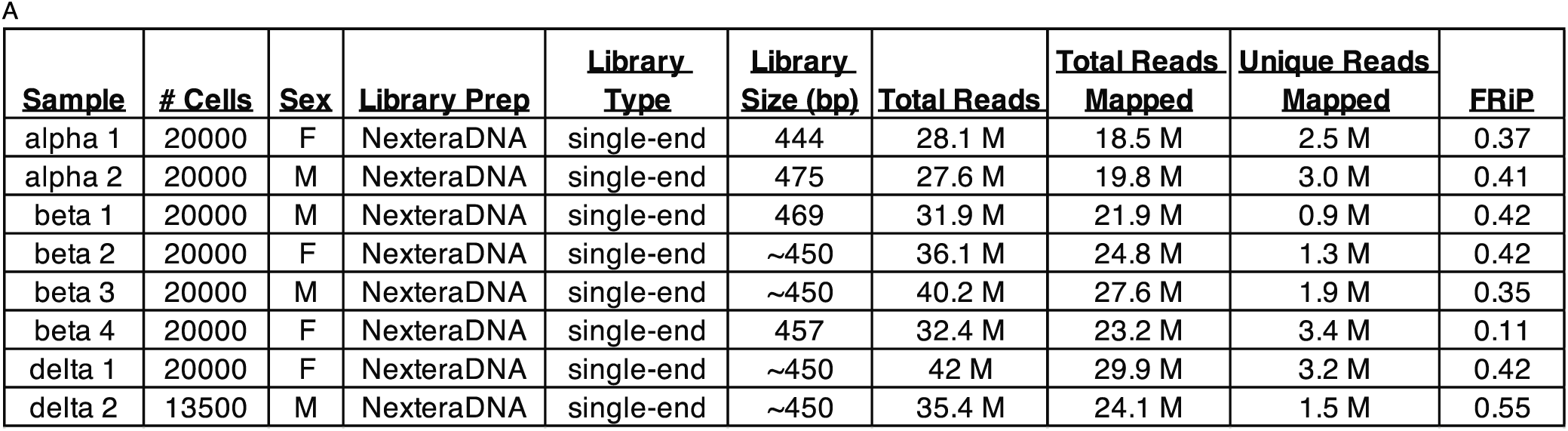
Quality control metrics across all ATAC-Seq replicates described.

**Supplemental Table 2.** Aggregated dataset description and reference. A: Pancreatic islet ChIP Seq transcription factor data aggregated to identify enhancer and enhancer regions. B: Pancreatic islet histone data aggregated to identify enhancer and enhancer regions. The final approach utilized two histone marks deemed most relevant at delineating putative enhancer regions while taking into account a risk of both false positives and false negatives.

| **ChIP (Transcription Factor) Datasets** |  |
| --- | --- |
| **<u>Marker</u>** | **<u>Accession Info</u>** |
| MafA | GSE30298 |
| Nkx2.2 | GSE79785 |
| Rfx6 | GSE62844 |
| Neurod1 | GSE54046 |
| Foxa2 | GSE54046 |
| Isl1 | GSE84759 |
| Kat2b | GSE78860 |
| Ldb1 | GSE84759 |
| Nkx6.1 | GSE40975 |
| Pdx1 | E-MTAB-1143 |
| Gata6 | GSE57090 |
| Insm1 | GSE54046 |

B
| **ChIP (Histone) Datasets** |  |
| --- | --- |
| **<u>Marker</u>** | **<u>Accession Info</u>** |
| H3k27ac | GSE110648 |
| H3k4me3 | GSE110648 |
| H3k4me1 | GSE68618 |
| H2ak119ub | GSE110648 |
| H3k27me1 | GSE110648 |
| H3k36me3 | GSE110648 |
| H3K9me3 | GSE110648 |
| H3k9ac | GSE87530 |
| H3k27me3 | GSE110648 |

**Supplemental Table 3.**
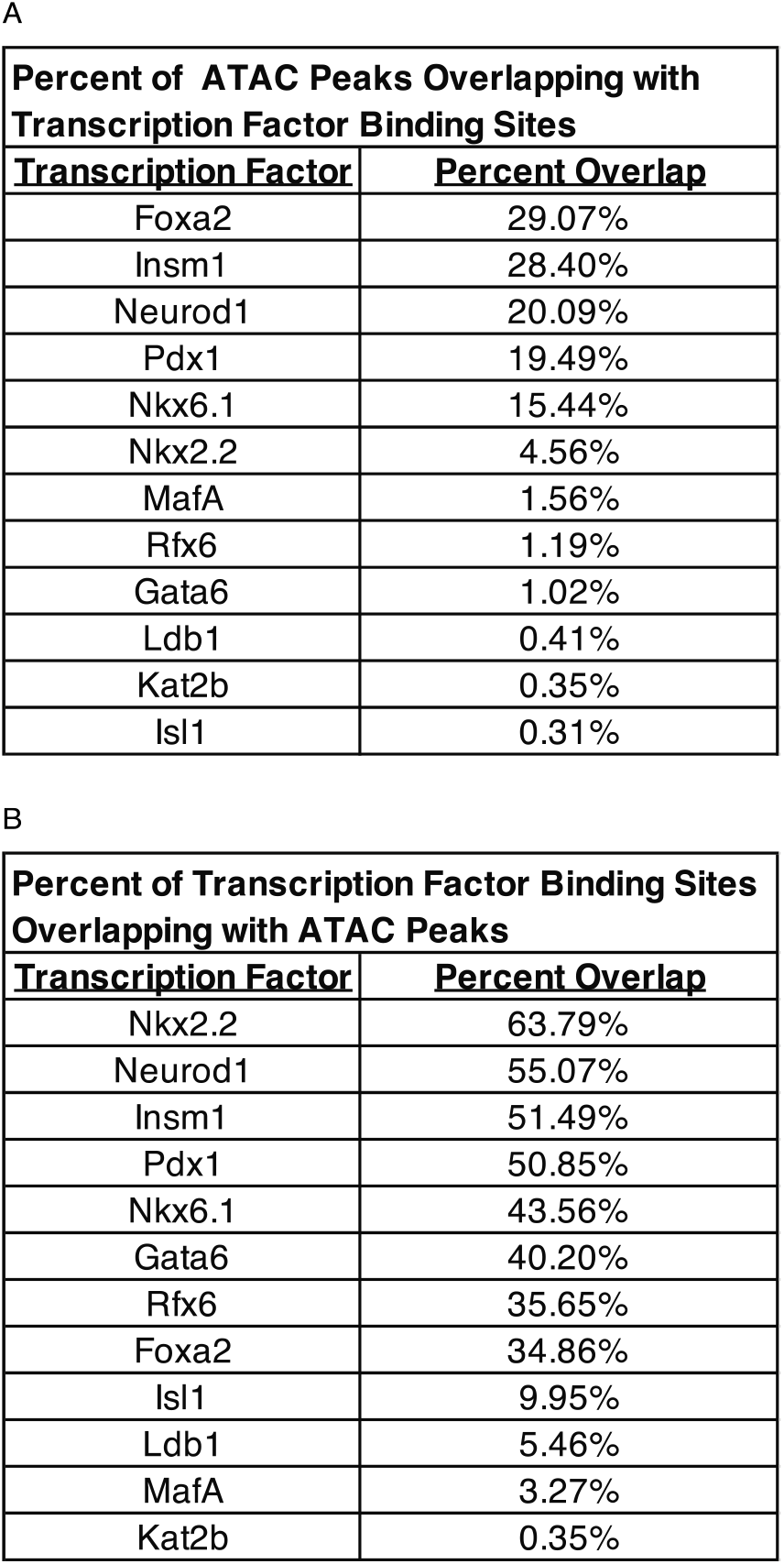
Validating open chromatin peaks against known pancreatic islet ChIP binding sites. A: Evaluating the extent of open chromatin– as defined by our ATAC-Seq consensus peak set – contained binding sites for known, pancreatic islet transcription factors. Percent of open chromatin with associated binding sites ranged from 0.31-29.07%. The transcription factors Foxa2, Insm1, and Neurod1 had the highest number of binding sites. B: Evaluating the extent of each ChIP-Seq experiment’s binding site calls overlapped with open chromatin. Percent of overlap ranged from 0.35-63.79%. Nkx2.2, Neurod1, and Insm1 had the greatest overlap.

| **Percent of ATAC Peaks Overlapping with Transcription Factor Binding Sites** |  |
| --- | --- |
| **Transcription Factor** | **Percent Overlap** |
| Foxa2 | 29.07% |
| Insm1 | 28.40% |
| Neurod1 | 20.09% |
| Pdx1 | 19.49% |
| Nkx6.1 | 15.44% |
| Nkx2.2 | 4.56% |
| MafA | 1.56% |
| Rfx6 | 1.19% |
| Gata6 | 1.02% |
| Ldb1 | 0.41% |
| Kat2b | 0.35% |
| Isl1 | 0.31% |

B
| **Percent of Transcription Factor Binding Sites Overlapping with ATAC Peaks** |  |
| --- | --- |
| **Transcription Factor** | **Percent Overlap** |
| Nkx2.2 | 63.79% |
| Neurod1 | 55.07% |
| Insm1 | 51.49% |
| Pdx1 | 50.85% |
| Nkx6.1 | 43.56% |
| Gata6 | 40.20% |
| Rfx6 | 35.65% |
| Foxa2 | 34.86% |
| Isl1 | 9.95% |
| Ldb1 | 5.46% |
| MafA | 3.27% |
| Kat2b | 0.35% |

**Supplemental Table 4.** Validating motif-calling approach against known ChIP binding sites. A: Pancreatic islet ChIP Seq transcription factor peak calls analyzed by the motif-calling method to determine sensitivity and specificity. True positive calls ranged from 0.59-57%, and false positives ranged from 1.19- 8.34%. B: Pancreatic islet ChIP Seq transcription factor peak calls limited to open chromatin determined by the consensus peak set analyzed by the motif-calling method to determine sensitivity and specificity. True positive calls ranged from 4.71-65.10%, and false positives ranged from 1.68-8.81%.

| **Transcription Factor Motif Calls against ChIP Peaks Validation** |  |  |  |  |
| --- | --- | --- | --- | --- |
| **Transcription Factor** | **False Negative** | **True Positive** | **False Positive** | **True Negative** |
| Rfx6 | 43.00% | 57% | 8.34% | 91.66% |
| Gata6 | 77.58% | 22.42% | 1.78% | 98.22% |
| Foxa2 | 78.17% | 21.83% | 6.38% | 93.62% |
| Nkx6.1 | 82.05% | 17.95% | 6.89% | 93.11% |
| Nkx2.2 | 89.48% | 10.52% | 2.98% | 97.02% |
| Insm1 | 93.39% | 6.61% | 3.63% | 96.37% |
| Pdx1 | 95.51% | 4.49% | 1.26% | 98.74% |
| Isl1 | 98.33% | 1.67% | 0.51% | 99.49% |
| MafA | 99.41% | 0.59% | 1.19% | 98.81% |

B
| **Transcription Factor Motif Calls against ChIP Peaks (Open ATAC only) Validation** |  |  |  |  |
| --- | --- | --- | --- | --- |
| **Transcription Factor** | **False Negative** | **True Positive** | **False Positive** | **True Negative** |
| Rfx6 | 34.90% | 65.10% | 8.81% | 91.19% |
| Foxa2 | 80.11% | 19.89% | 8.92% | 91.08% |
| Gata6 | 83.80% | 16.20% | 1.98% | 98.02% |
| Nkx6.1 | 88.61% | 11.39% | 8.74% | 91.26% |
| Nkx2.2 | 90.28% | 9.72% | 3.69% | 96.31% |
| MafA | 91.29% | 8.71% | 1.73% | 98.27% |
| Insm1 | 91.74% | 8.26% | 3.97% | 96.03% |
| Isl1 | 94.12% | 5.88% | 0.74% | 99.26% |
| Pdx1 | 95.30% | 4.71% | 1.68% | 98.32% |

## Supplemental Figures

**Supplemental Figure 1.**
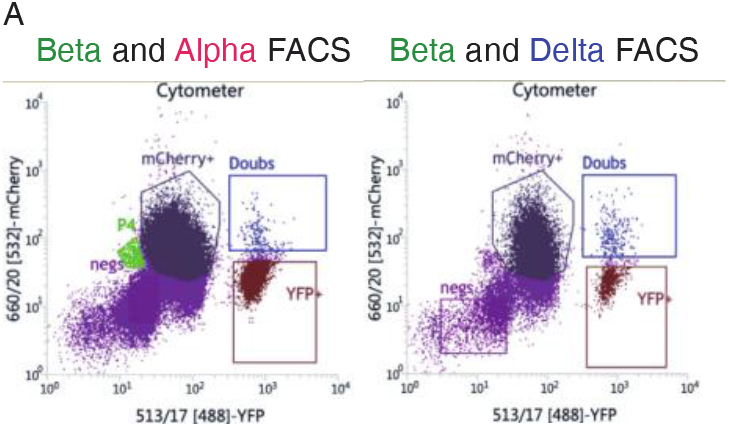
FACS sorting gates used to isolate alpha, beta, and delta cells through our mouse reporter lines. FACS sorting gates isolating beta cells (Ins2-mCherry+) from either alpha (Gcg- YFP+) or delta cells (Sst-YFP+). Double negatives are non-beta and non-alpha or non-delta cells. Double positives (mCherry/YFP+) represent cells with both Ins2 expression and Gcg or Sst expression, reflective of transdifferentiated beta cells. These were not included in any of the samples.

**Supplemental Figure 2.**
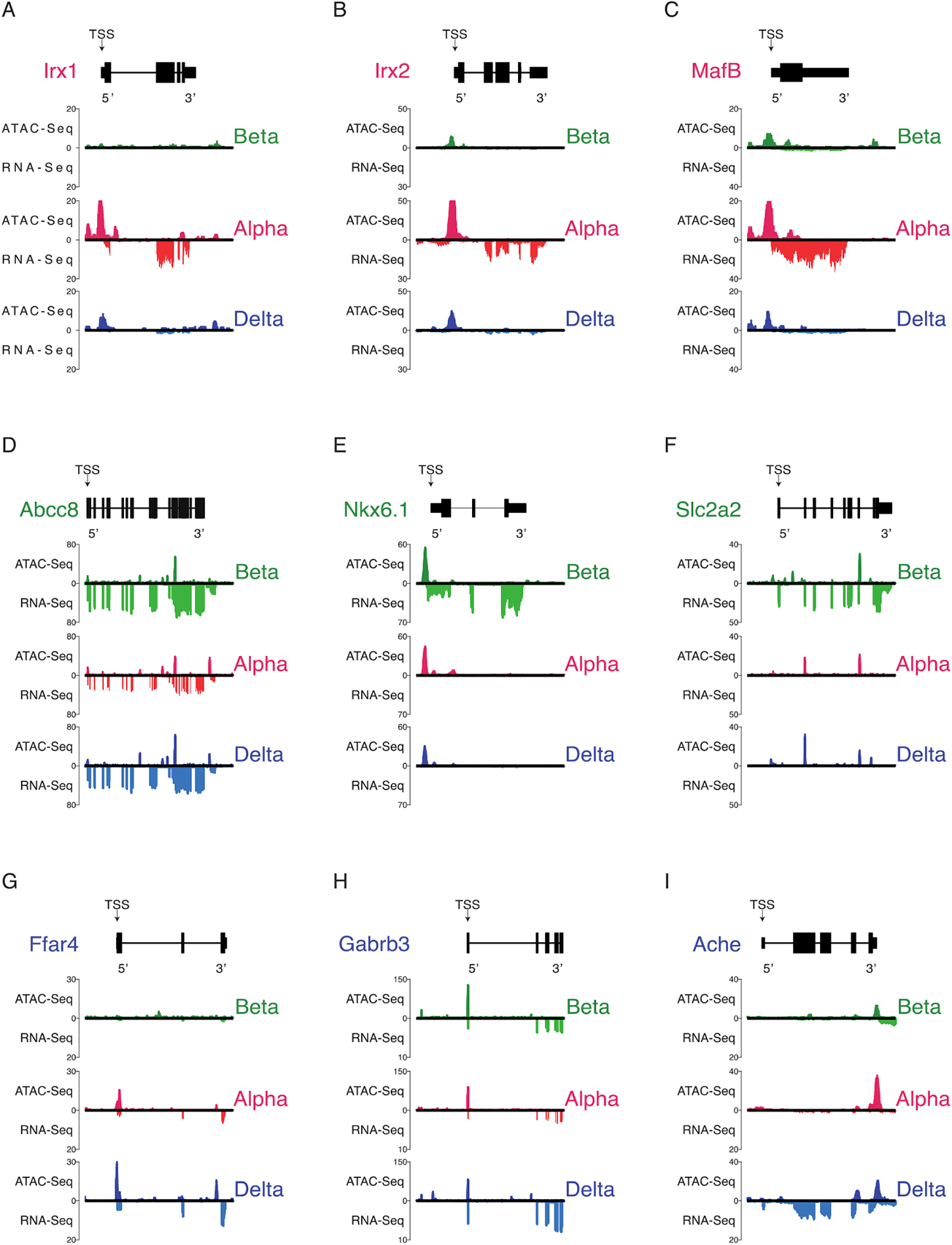
Validating more chromatin accessibility ATAC Seq and companion RNA-Seq expression in alpha, beta, and delta cells against hallmark genes governing its respective cell’s identity. All genes are oriented for 5’ to 3’ end. A-C: Chromatin accessibility and transcript expression across alpha cell hallmark genes *Irx1, Irx2,* and *MafB*. D-F: Chromatin accessibility and transcript expression across beta cell hallmark genes *Abcc8, Nkx6.1,* and *Slc2a2.* G-I: Chromatin accessibility and transcript expression across delta cell hallmark genes *Ffar4, Gabrb3,* and *Ache*.

**Supplemental Figure 3.**
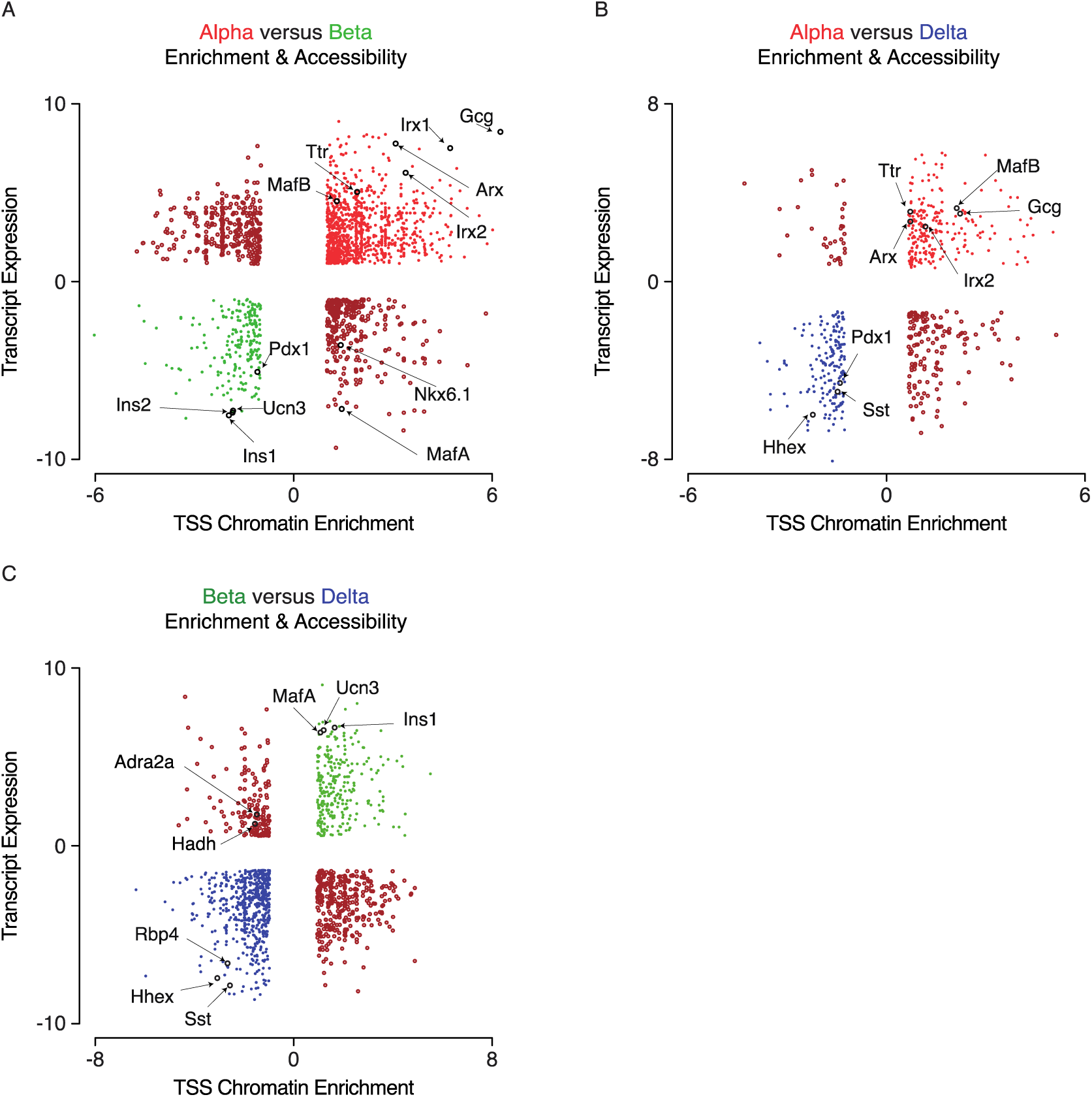
Chromatin enrichment does not always correlate with associated gene expression. Select hallmark genes defining demonstrating congruent and incongruent chromatin and gene enrichment for cell-specific markers. A: Differentially enriched chromatin at TSS regions and respective gene expression between alpha and beta cells. The majority of cell-specific markers show TSS- enrichment within the cell type of expression. Notably, *Nkx6.1* and *MafA* show TSS enrichment in alpha cells, despite being transcription factors associated with beta cells. B: Differentially enriched chromatin at TSS regions and respective gene expression between alpha and delta cells. C: Differentially enriched chromatin at TSS regions and respective gene expression between beta and beta cells. The majority of cell-specific markers show TSS-enrichment within the cell type of expression.

**Supplemental Figure 4.**
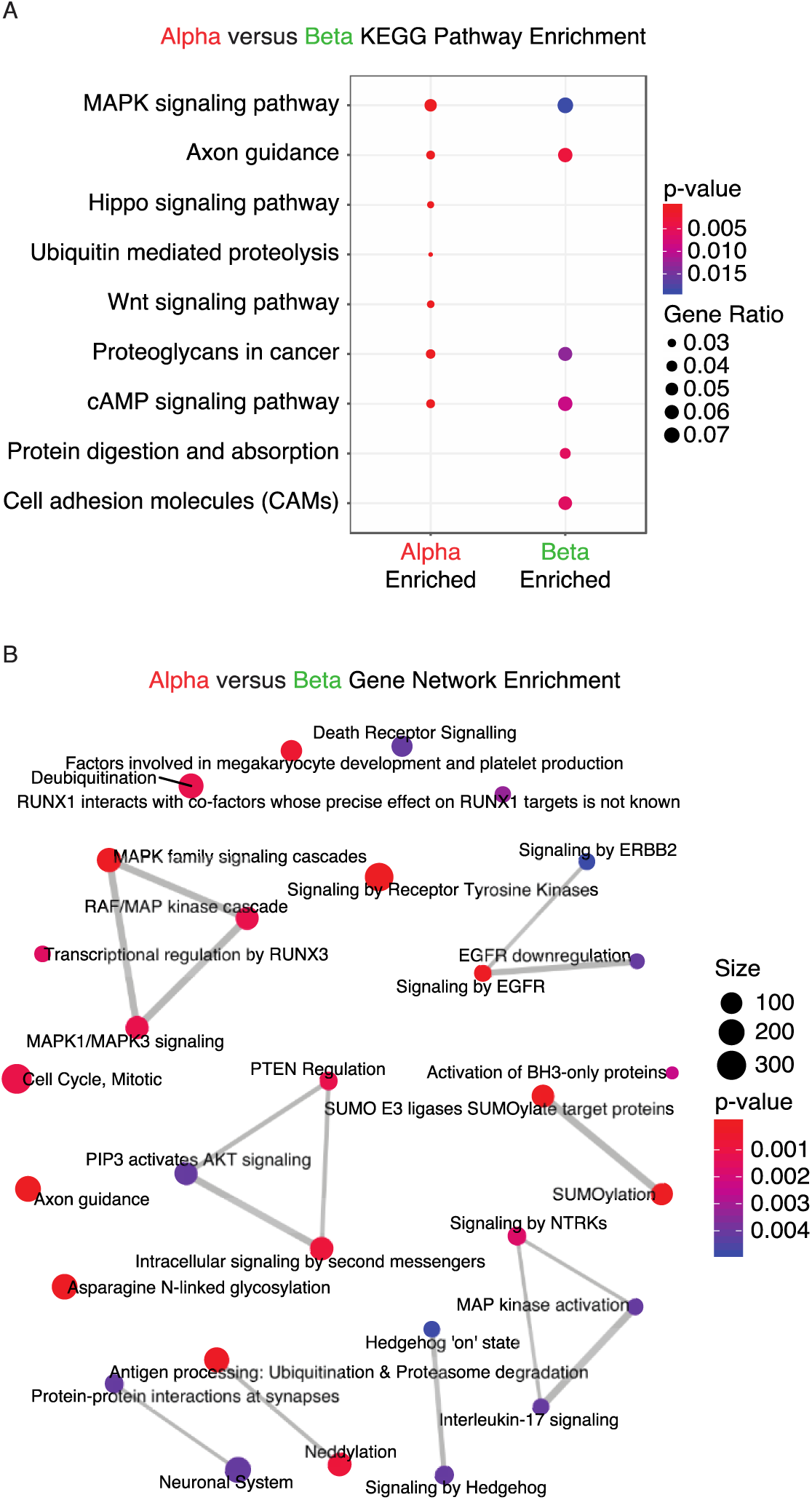
Evaluating KEGG and gene network enrichment across differentially enriched peaks between alpha and beta cells. A: KEGG enrichment of differentially enriched peaks identified pathways common between the two cell types, or unique to one. B: Gene network enrichment indicative of possible functions of differentially enriched chromatin regions between the two cell types.

**Supplemental Figure 5.**
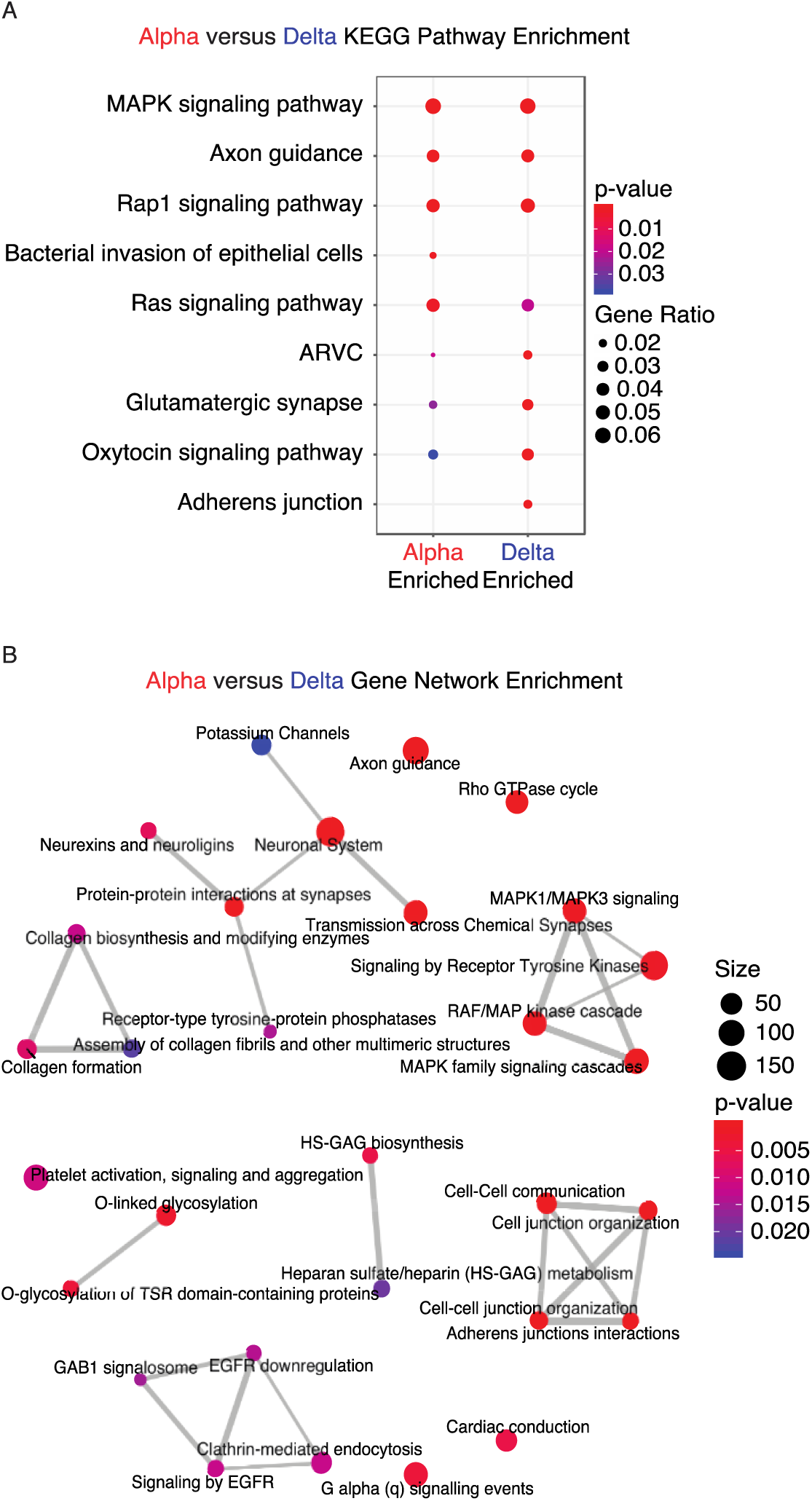
Evaluating KEGG and gene network enrichment across differentially enriched peaks between alpha and delta cells. A: KEGG enrichment of differentially enriched peaks identified pathways common between the two cell types, or unique to one. B: Gene network enrichment indicative of possible functions of differentially enriched chromatin regions between the two cell types.

**Supplemental Figure 6.**
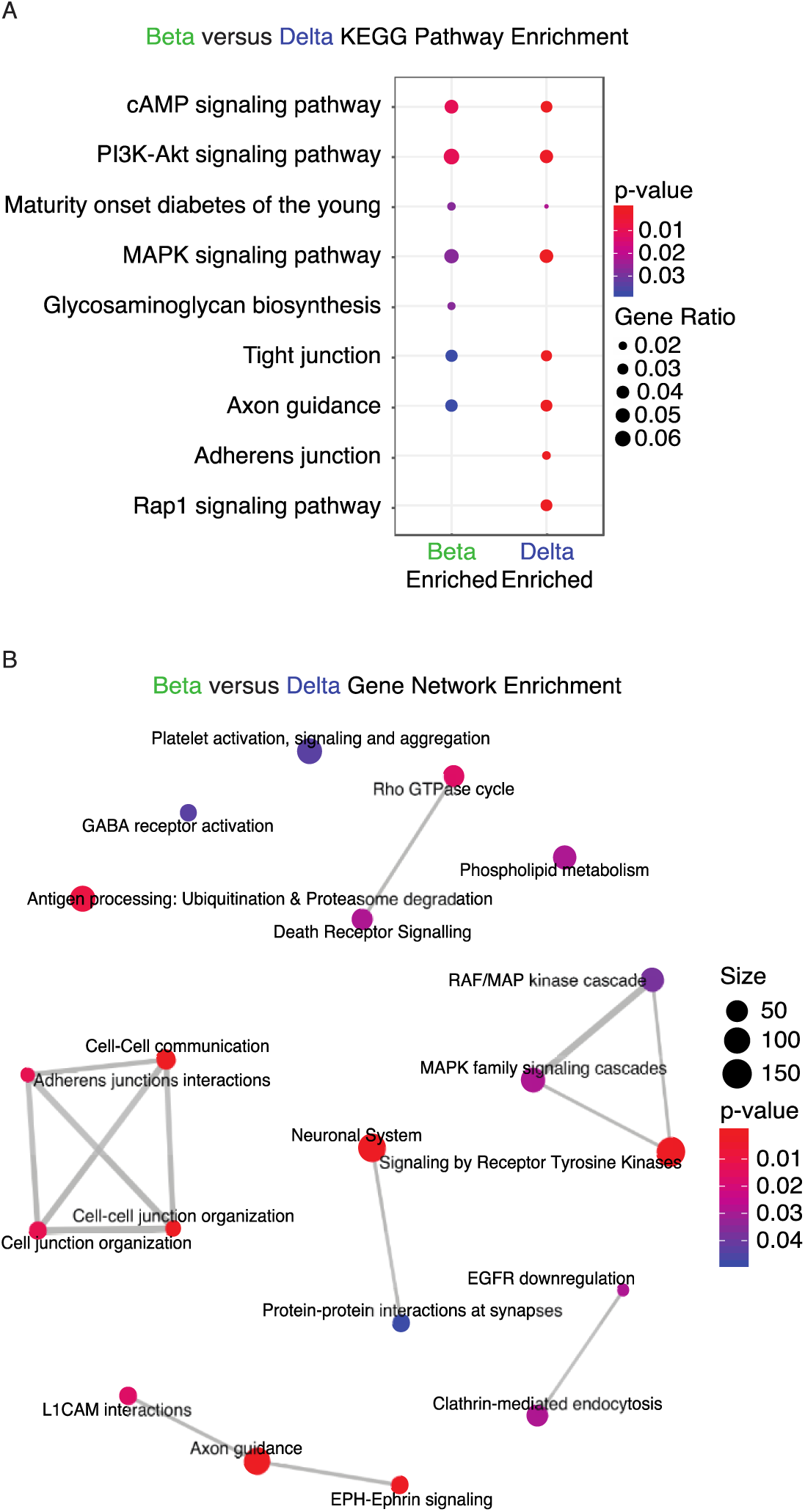
Evaluating KEGG and gene network enrichment across differentially enriched peaks between beta and delta cells. A: KEGG enrichment of differentially enriched peaks identified pathways common between the two cell types, or unique to one. B: Gene network enrichment indicative of possible functions of differentially enriched chromatin regions between the two cell types.

**Supplemental Figure 7.**
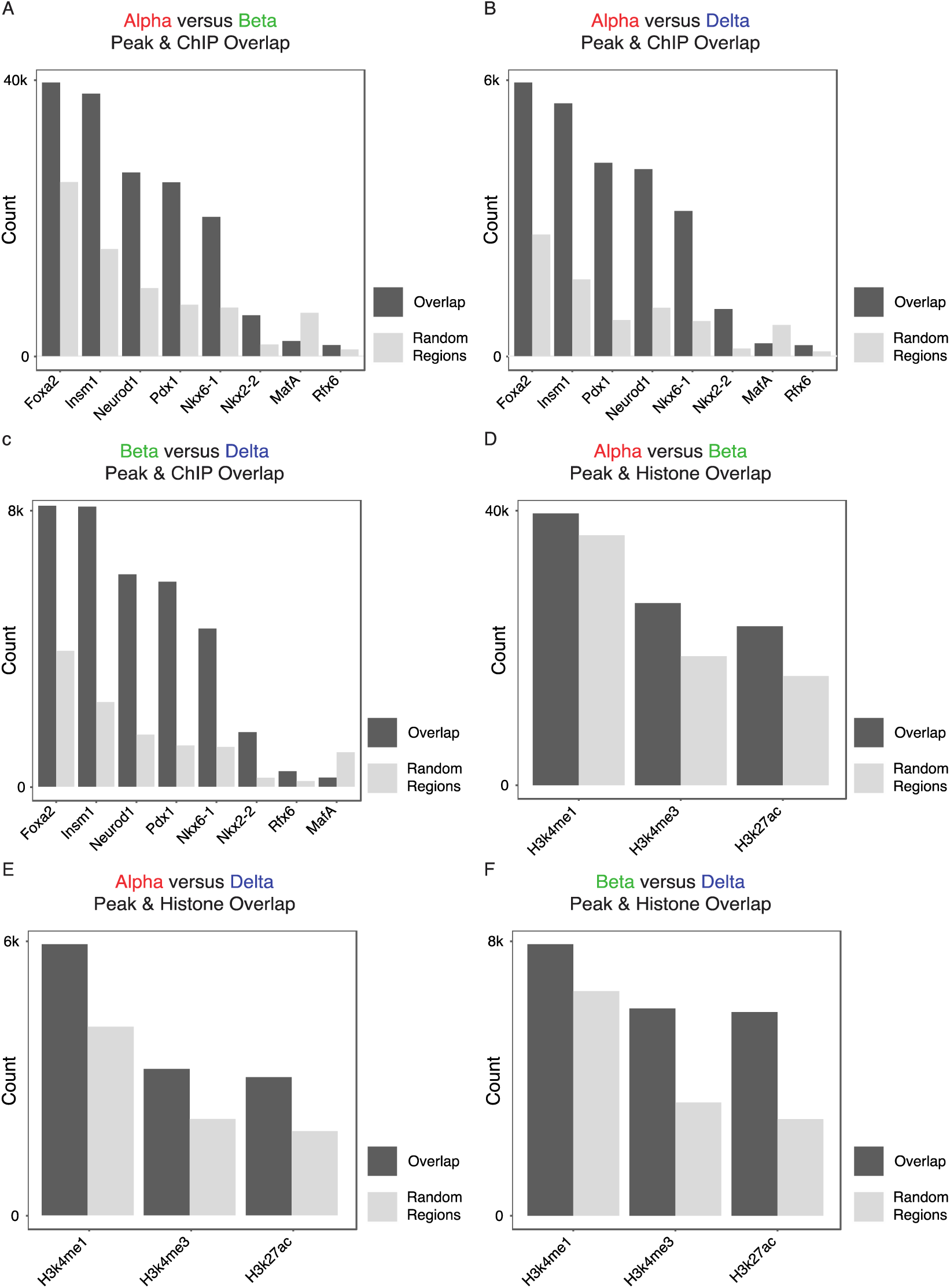
Verifying transcription factor binding sites and histone mark occurrence at chromatin peaks to determine significance (observed versus expected). A-C: Transcription factors on chromatin regions deemed enriched between differentially enriched chromatin across all three pairwise comparisons.. The majority of transcription factors used in our analysis were deemed statistically significant when observed compared to predicted. D-F: Histone mark occurrence on chromatin regions deemed enriched between differentially enriched chromatin across all three pairwise comparisons. All histone marks used in our analysis were deemed statistically significant when observed compared to predicted.

**Supplemental Figure 8.**
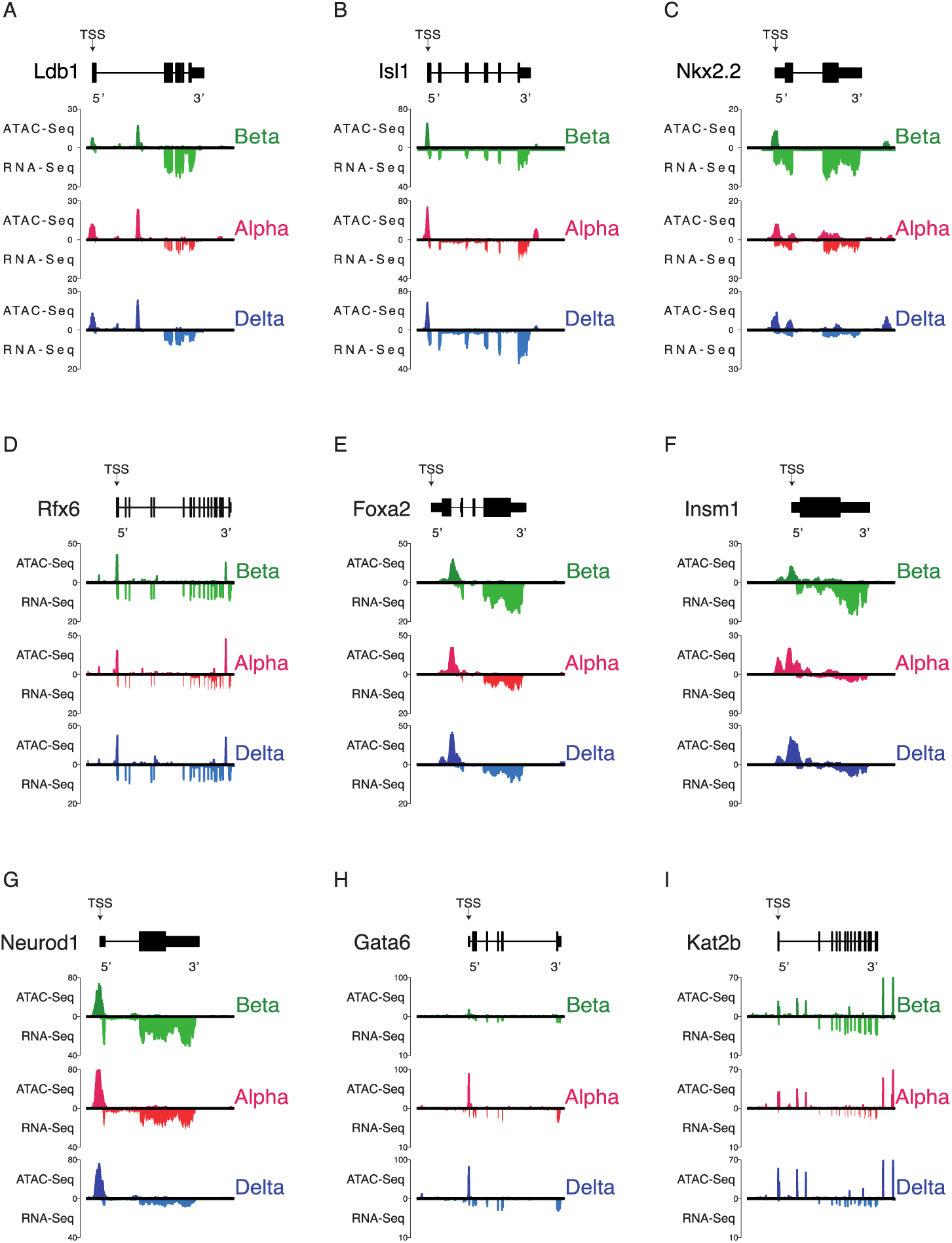
Aggregated transcription factor ATAC Seq and companion RNA-Seq expression in alpha, beta, and delta cells. All genes are oriented for 5’ to 3’ end. A-I: Chromatin accessibility and gene expression for aggregated ChIP datasets.

**Supplemental Figure 9.**
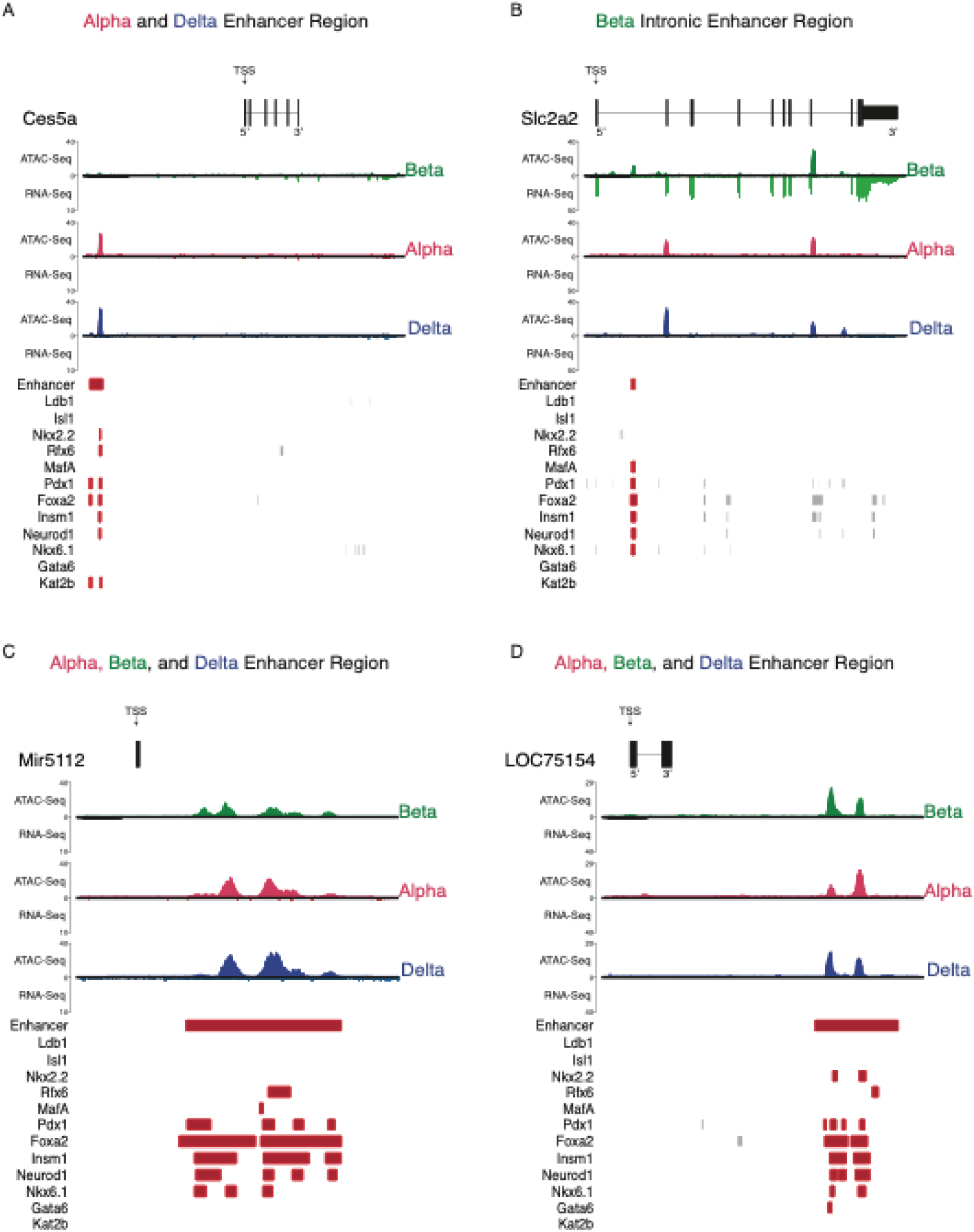
Further illustration of enhancer calls. A: Visualizing a common alpha and delta enhancer region, unavailable in beta cells. B: Further illustration of a beta-unique enhancer region, occurring on the first intron of *Slc2a2,* with 6 co-binding sites for multiple transcription actors. C-D: Two examples of called enhancer regions common across all three cell types. Both are in distal-intergenic regions of the genome and exhibit high transcription factor co-binding activity.

## Supplemental Datasets

**Supplemental Dataset 1 – Annotated consensus chromatin peak set across alpha, beta and delta cells, along with differential enrichment results between the three pairwise comparisons.**

**Supplemental Dataset 2 – Congruent and incongruent genes of differentially expressed genes between the three pairwise comparisons.** Congruent genes showed gene expression in the same direction as chromatin accessibility enrichment, whereas incongruent genes had opposing expression and enrichment.

**Supplemental Dataset 3 – Unfiltered putative enhancer calls defined by open chromatin region in at least one of three cell types, overlapping the histone markers H3K27ac and H3K4me1.** These regions were crossed against pancreatic islet transcription factors to identify enhancer regions.

**Supplemental Dataset 4 – Filtered putative enhancer calls defined by open chromatin region in at least one of three cell types, overlapping the histone markers H3K27ac and H3K4me1.** These regions were crossed against pancreatic islet transcription factors to identify enhancer regions. Last, these data were filtered for regions occurring on curated enhancer calls in the mouse genome using the FANTOM5 database.

